# Increasing hub disruption parallels dementia severity in autosomal dominant Alzheimer disease

**DOI:** 10.1101/2023.10.29.564633

**Authors:** Jiaxin Cindy Tu, Peter R. Millar, Jeremy F. Strain, Andrew Eck, Babatunde Adeyemo, Alisha Daniels, Celeste Karch, Edward D. Huey, Eric McDade, Gregory S. Day, Igor Yakushev, Jason Hassenstab, John Morris, Jorge J. Llibre-Guerra, Laura Ibanez, Mathias Jucker, Patricio Chrem Mendez, Randell J. Bateman, Richard J. Perrin, Tammie Benzinger, Clifford R. Jack, Richard Betzel, Beau M. Ances, Adam T. Eggebrecht, Brian A. Gordon, Muriah D. Wheelock, the Dominantly Inherited Alzheimer Network

## Abstract

Hub regions in the brain, recognized for their roles in ensuring efficient information transfer, are vulnerable to pathological alterations in neurodegenerative conditions, including Alzheimer Disease (AD). Given their essential role in neural communication, disruptions to these hubs have profound implications for overall brain network integrity and functionality. Hub disruption, or targeted impairment of functional connectivity at the hubs, is recognized in AD patients. Computational models paired with evidence from animal experiments hint at a mechanistic explanation, suggesting that these hubs may be preferentially targeted in neurodegeneration, due to their high neuronal activity levels—a phenomenon termed “activity-dependent degeneration”. Yet, two critical issues were unresolved. First, past research hasn’t definitively shown whether hub regions face a higher likelihood of impairment (targeted attack) compared to other regions or if impairment likelihood is uniformly distributed (random attack). Second, human studies offering support for activity-dependent explanations remain scarce.

We applied a refined hub disruption index to determine the presence of targeted attacks in AD. Furthermore, we explored potential evidence for activity-dependent degeneration by evaluating if hub vulnerability is better explained by global connectivity or connectivity variations across functional systems, as well as comparing its timing relative to amyloid beta deposition in the brain. Our unique cohort of participants with autosomal dominant Alzheimer Disease (ADAD) allowed us to probe into the preclinical stages of AD to determine the hub disruption timeline in relation to expected symptom emergence.

Our findings reveal a hub disruption pattern in ADAD aligned with targeted attacks, detectable even in pre-clinical stages. Notably, the disruption’s severity amplified alongside symptomatic progression. Moreover, since excessive local neuronal activity has been shown to increase amyloid deposition and high connectivity regions show high level of neuronal activity, our observation that hub disruption was primarily tied to regional differences in global connectivity and sequentially followed changes observed in Aβ PET cortical markers is consistent with the activity-dependent degeneration model. Intriguingly, these disruptions were discernible 8 years before the expected age of symptom onset.

Taken together, our findings not only align with the targeted attack on hubs model but also suggest that activity-dependent degeneration might be the cause of hub vulnerability. This deepened understanding could be instrumental in refining diagnostic techniques and developing targeted therapeutic strategies for AD in the future.

## Introduction

Alzheimer disease (AD) is the most common neurodegenerative disease that manifests as progressive loss of cognitive functions and affects millions of people worldwide. It is characterized by a cascade of complex pathologic changes in the brain including amyloid beta aggregation and tau tangles, resulting in neurodegeneration^1^. While these microscopic changes have been well-documented, there is growing interest in understanding how these pathologies translate to altered brain connectivity patterns observed in AD patients.

Resting-state functional connectivity (FC), as measured with temporal correlations of the blood oxygen level-dependent (BOLD) signals between regions of the brain from fMRI data collected at a task-free state^2^, differs in clinical and pre-clinical AD individuals versus cognitively normal controls^3–7^. FC is a widely accessible and non-invasive tool for assessing brain organization, and a potential imaging marker for AD^8–11^ since disruptions in connectivity between brain regions may be linked to synaptic changes before cell death and atrophy^12^. Prospective imaging studies suggested that the posterior parts of the default mode network deteriorate earlier than anterior parts in AD, providing evidence for a cascading network failure mechanism^13,14^. Importantly, the initiation of amyloid^15,16^, and tau^17^ pathologies as well as the rate of their accumulation^18,19^ in the brain is not spatially homogenous, providing converging evidence for differences in regional vulnerability to pathological changes. However, precisely what underlies the sequence of regional FC failure, as well as how FC disruptions relate to the molecular pathology is unknown.

Hub regions^20^ are highly connected nodes with high network centrality that play a critical role in facilitating efficient communication and integration of information across different regions of complex networks. Brain hubs are affected across multiple diseases^21^ including AD^22,23^. One hypothesis for this hub vulnerability to pathology and degeneration is that hub regions are selectively targeted by activity-dependent damage^24^. Several lines of evidence support this hypothesis. 1) Hubs have high metabolic demands^12,25–27^; 2) they are especially susceptible to amyloid beta deposition in AD^12,27–29^; and, 3) they serve as the spreading centers for tau pathology^17,30,31^. Due to their topologically central role, targeted attacks on hubs have a more deleterious effect on network efficiency^21,32–34^. Indeed, FC alterations especially at cortical hubs^12,23^ have been identified in AD patients compared to cognitively normal controls in previous research as well as in mice with extracellular amyloidosis (TgCRND8 mice)^35^. *In vivo* studies in mice validate the relationship between neuronal activity level and amyloid-beta deposition^36,37^, suggesting that increasing amyloid burden through increased baseline activity triggers hub disruptions. While there is strong theoretical and empirical evidence to support the role of hub disruption in AD, little is known concerning the relationship between hub disruption, dementia severity, and symptomatic onset.

Here we leverage a unique population with autosomal dominant AD (ADAD), which allows for accurate estimation of years until symptom onset (EYO) due to the highly predictable onset of cognitive decline^38^. As a genetic form of the disease, our ADAD participants have a high certainty in AD diagnosis (as opposed to other forms of dementias). Furthermore, since ADAD occurs in a younger population (<60 years) than sporadic AD, we can test associations with AD pathology with minimal confounding co-pathology and can estimate FC network characteristics with minimal age-related changes in neurovascular coupling^39^.

We examine the regional vulnerability in terms of lower FC by measuring FC hub disruption as a function of ADAD dementia progression from pre-clinical (Clinical Dementia Rating® [CDR®] =0) to moderate and severe dementia (CDR≥1). We hypothesize that hub disruption is an early-emerging phenomenon that intensifies with disease progression. Finally, we investigated the relative timing of hub disruption compared to cortical amyloid deposition and cognitive decline. Our goal is to test the targeted attack on hubs model in ADAD over the course of disease progression and obtain evidence for activity-dependent degeneration.

## Materials and methods

### Participants

Individuals were recruited from the Dominantly Inherited Alzheimer Network (DIAN) Observational Study (https://dian.wustl.edu/). Here we examine cross-sectional data from mutation carriers (MC; N = 122) and unaffected non-carriers (NC; N = 85) family members (Supplementary Table 1) with alterations in presenilin 1 (PSEN1), presenilin 2 (PSEN2), or the amyloid precursor proteins (APP)^38^. Age at symptom onset is relatively consistent within families and mutation types; this allows participants to be staged by their EYO^38,40^. *Both MCs and NCs have an EYO value based their familial pedigree but only MCs are expected to develop ADAD*. The study was reviewed and approved by the institutional review board at Washington University in St. Louis and written informed consent forms obtained from participants or their legally authorized representatives in accordance with their local institutional review board. The data are from the 15^th^ annual data freeze.

### CDR stages

The MCs were further staged by dementia severity using the global Clinical Dementia Rating (CDR(Morris, 1993) into three groups: cognitively normal (CDR = 0), very mild dementia (CDR = 0.5), and mild-to-severe dementia (CDR>=1). To control for the effect of aging, we age-matched the NCs for each MC group according to the following procedure. First, Z-scores were calculated for the age and EYO values separately using their mean and standard deviation across all participants, resulting in a vector of size 2×1 for each participant. The Euclidean distances between the vectors were calculated and the closest MC for each NC participant was determined, defining an age-matched group of NCs for each MC group (Table 1).

**Table 1.**
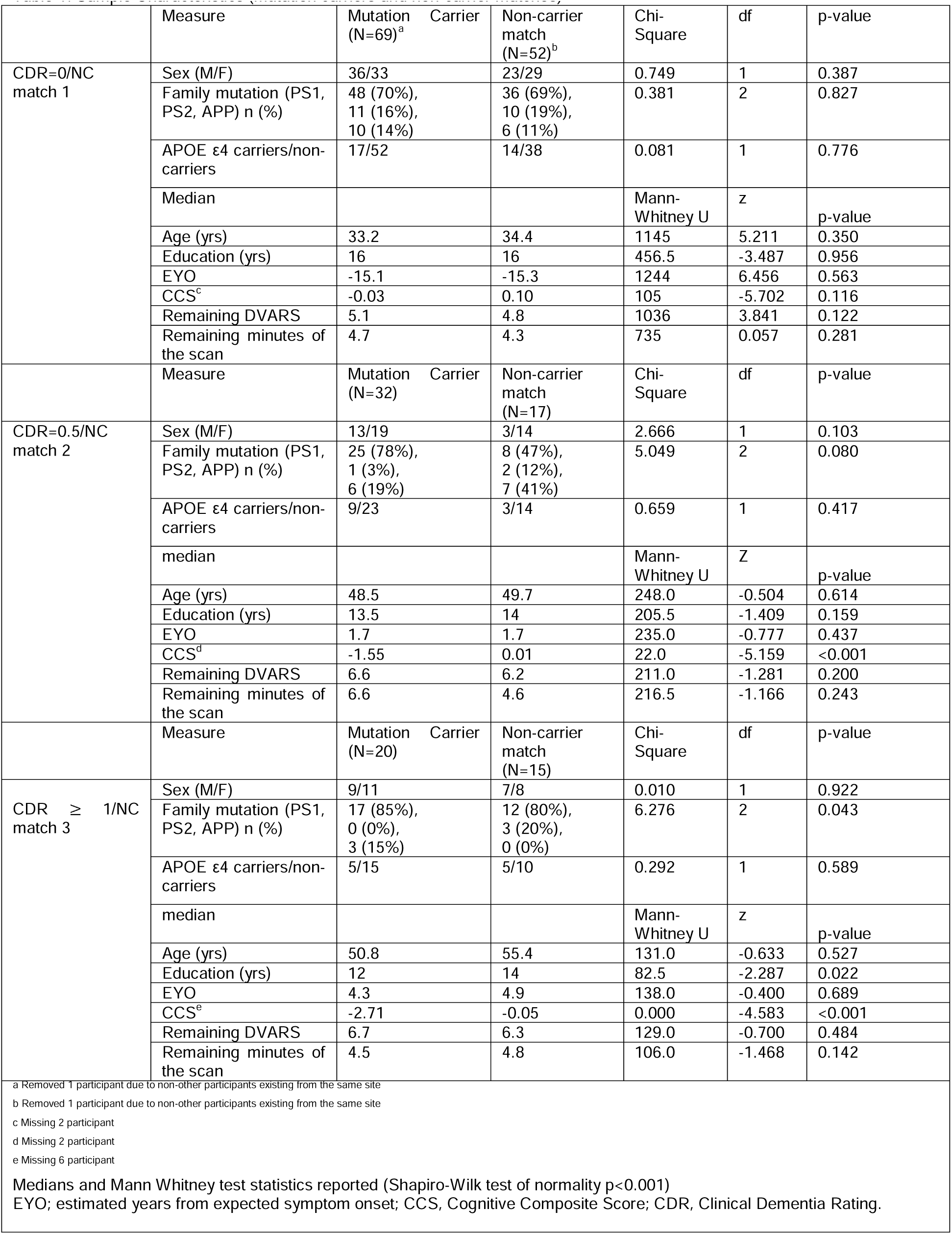
Sample Characteristics (mutation carriers and non-carrier matches)

### MRI Data Acquisition

Neuroimaging protocols have been previously published^40^. Briefly, T1-weighted magnetization-prepared rapid acquisition gradient echo (MP-RAGE) images were acquired at multiple sites on Siemens 3T scanners (Erlangen, Germany). Resting-state fMRI scans were acquired with echo planar imaging (EPI) while participants were instructed to maintain visual fixation on a crosshair. The sequence details are provided in the Supplementary Materials (Supplementary Table 2). “Pre-scan normalize” was enabled to minimize gain field inhomogeneities attributable to proximity to the receiver coils. Acquisition lasted ∼6 minutes each run and the number of acquired runs in the DIAN cohort varied between 1 and 3.

### MRI Data Pre-processing

Details on pre-processing followed previously described methods^6,41^ using the 4dfp suite of tools (http://4dfp.readthedocs.io). Briefly, slice timing correction and intensity normalization were performed. Head motion was corrected within and across runs. The initial atlas transformation was computed by affine registration of the functional MRI data to an atlas-representative template via the MP-RAGE (EPI_mean_ → MP-RAGE → template). A final atlas transformation was performed after denoising. Frames with high motion, as measured by DVARS (frame-to-frame signal change over the entire brain) and the frame displacement (FD) measures^42^, were censored. The DVARS criterion was individually set to accommodate baseline shifts (see Supporting Information in^43^) and the FD criterion was 0.4 mm. Frames were censored if either criterion was exceeded. The time series were band-pass filtered between 0.005 Hz and 0.1 Hz. Censored frames were approximated by linear interpolation for band-pass filtering only and excluded from subsequent steps.

Denoising was then performed with a CompCor-like strategy^44^. As previously described^45^, nuisance regressors were derived from three compartments (white matter, ventricles, and extra-axial space) and were then dimensionality-reduced. White matter and ventricular masks were segmented in each participant using FreeSurfer 5.3^46^ and spatially resampled in register with FC data. The final set of nuisance regressors also included the six parameters derived from rigid body head-motion correction, the global signal averaged over the (FreeSurfer-segmented) brain, and the global signal temporal derivative. Finally, the volumetric time-series were non-linearly warped to Montreal Neurological Institute (MNI) 152 space (3mm)^3^ voxels using FNIRT^47^.

### Functional Connectivity

We selected 246 functional regions of interest (ROIs) separated into 13 networks throughout the cortical and subcortical areas as previously described^48^. The functional ROIs are a combination of cortical ROIs^49^ and subcortical ROIs^50^ (Figure 1A). Regions not reliably covered by the field of view (FOV) such as the cerebellar ROIs were excluded. A list of ROI coordinates and anatomical assignments has been described in previous publications^41,48^ and can be found in the Supplementary Materials (Supplementary Table 3). FC was estimated using zero-lag Pearson correlations calculated between 246 ROIs and Fisher-Z-transformed to improve normality. The resultant FC matrix can be represented as a graph with nodes as individual ROIs and edges with weights as the correlation values z(r). Group-average FC was generated by averaging the z(r) values across individual FC matrices within each of the CDR and age-matched NC groups (Figure 2).

**Figure 1.**
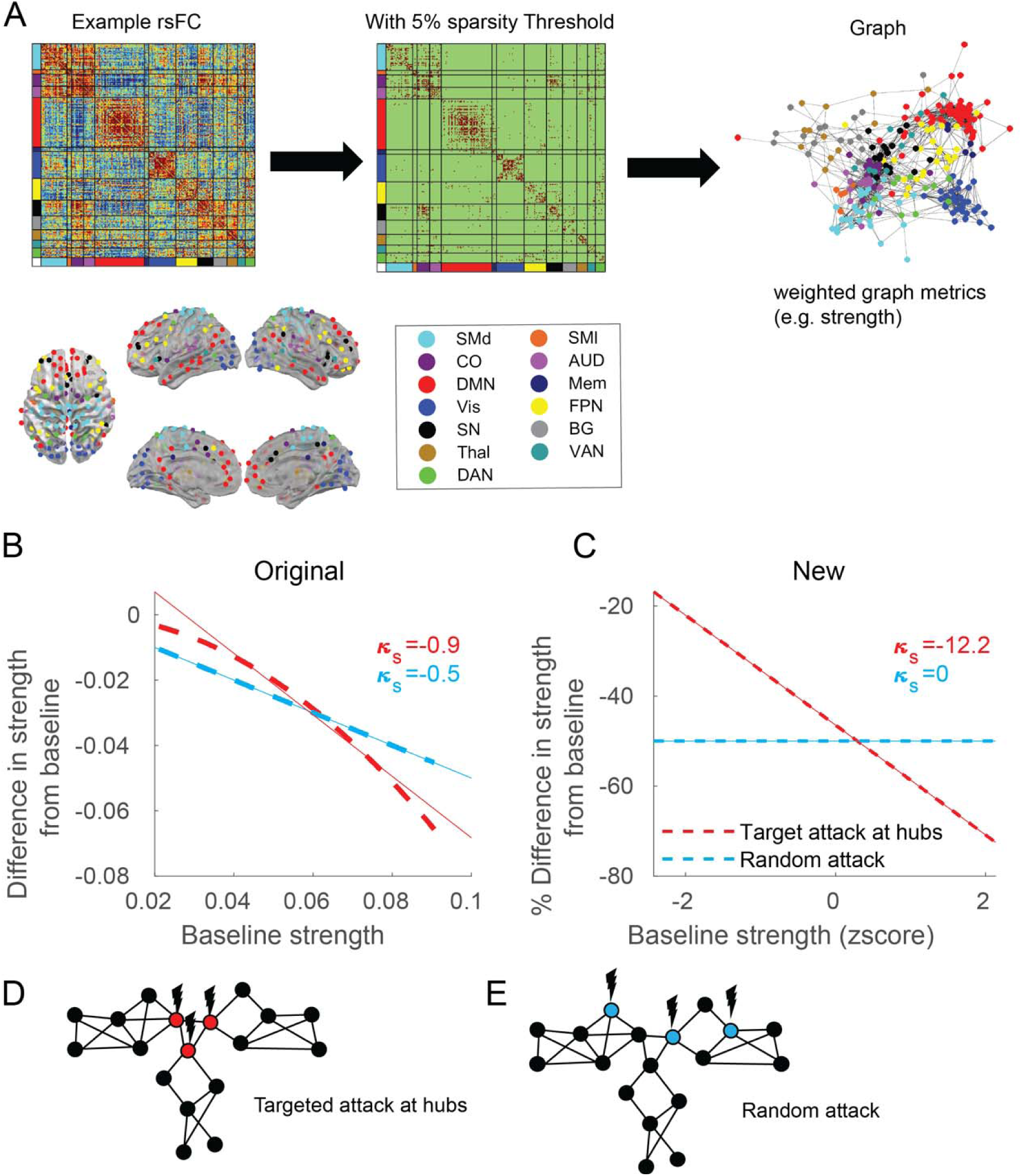
Graph theory method and hub disruption. **(A)** A resting-state functional connectivity (FC) is obtained from the Pearson correlation of the time series in each of the 246 cortical and subcortical pre-defined region of interest (ROI) pairs. The ROIs belong to 13 Networks: SMd, somatomotor dorsal; SMl, somatomotor lateral; CO, cingulo-opercular; AUD, auditory; DMN, default mode network; Mem, memory network; Vis, visual network; FPN, frontoparietal network; SN, salience network; BG, basal ganglia; Thal, thalamus; VAN, ventral attention network; DAN, dorsal attention network. Following convention in previous literature, a sparse graph is generated by thresholding the rsFC matrix at an edge density threshold of 5% starting from the maximum spanning tree (MST) backbone to ensure the connectedness of the graph. However, to demonstrate that our results are not limited to the threshold choice we also applied other thresholds. The graph generated has weighted edges that preserve the strength of individual connections. **(B)** Original method of hub disruption calculation. **(C)** New method of hub disruption calculation. **(D)** Cartoon illustration of targeted attack at the hubs. **(E)** Cartoon illustrating random damage.

**Figure 2.**
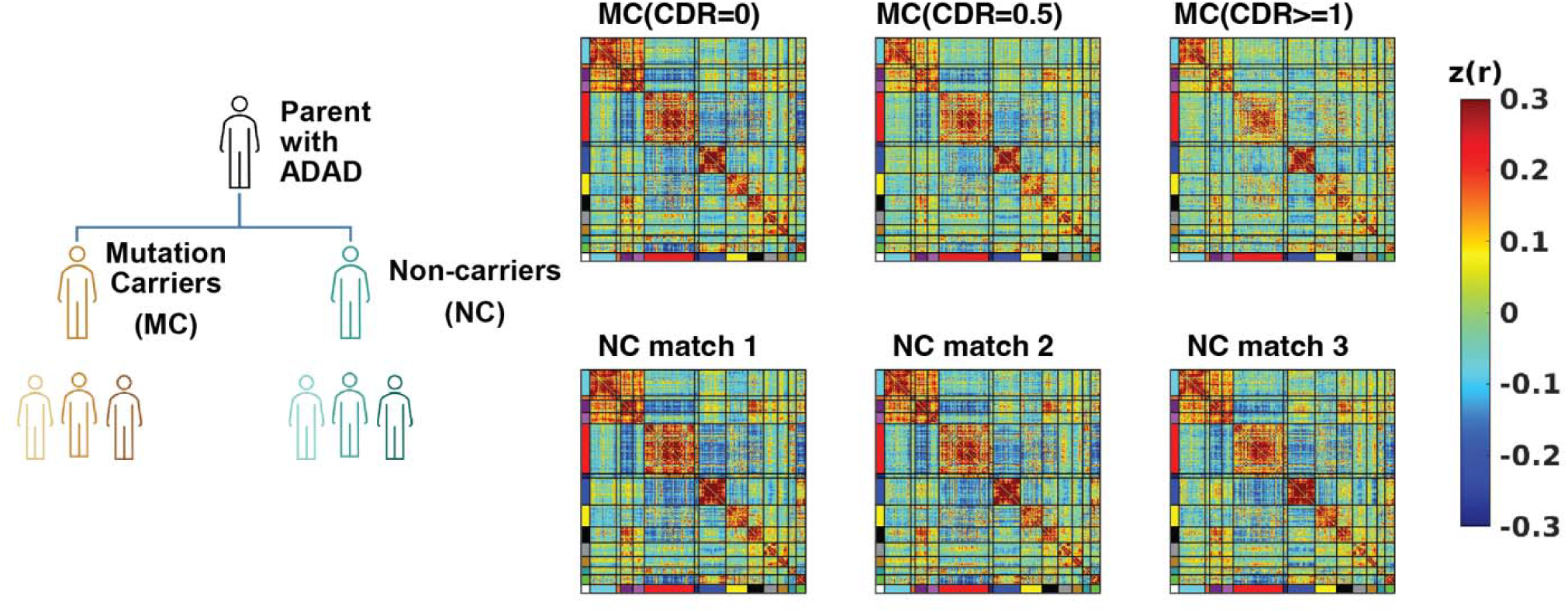
Functional connectivity (FC) within DIAN participant groups. Mean (lower-triangle) and standard deviation (upper-triangle) of Fisher Z-transformed FC matrix of 246 region of interests for mutation carriers (MC) at 3 Clinical Dementia Rating stages (CDR = 0, CDR = 0.5, CDR>=1) and corresponding age and EYO matched non-carrier (NC) groups. The FC is sorted by the networks in Figure 1 with corresponding colors.

### Data Harmonization

We used Correcting Covariance Batch Effects (CovBat, https://github.com/andy1764/CovBat_Harmonization)^51^ to remove site effects in mean, variance, and covariance on FC data, with age, mutation, EYO, education, CDR, sex, mutation gene type (PSEN1/PSEN2/APP), and APOE alleles included as the biological covariates that should be protected for during the removal of site effects. After CovBat, two participants (one MC and one NC) were removed from the analysis because they were only represented by a single site and harmonization could not be performed by the CovBat algorithm. The final sample size for analysis was MC = 121 and NC = 84. Similar qualitative results were obtained without the CovBat correction.

### Graph Theory Metrics

All graph theory metrics were calculated using the Brain Connectivity Toolbox (BCT, <u>brain-connectivity-toolbox.net</u>), a MATLAB toolbox for complex brain-network analysis^52^. Since regions with a high total positive connectivity tend to have a high total negative connectivity (Supplementary Figure 1), we asymmetrically weighed the positive and negative edges for all measures according to their relative magnitude at a given node following previous literature^53^. This allows for non-zero strengths at a full FC matrix without threshold (Supplementary Figure 2). Strength is calculated as the sum of signed edge weights around a node in a graph (Figure1E). This effectively measures global connectivity at an ROI. We also calculated two additional measures of centrality concerning module affiliations^54^: the within-module strength Z-Score (Z) and participation coefficient. Z measures how “well-connected” a node is to other nodes in the module. The participation coefficient measures the diversity of intermodular connections of individual nodes and within-module strength Z-score.

Given that there is no gold standard method for thresholding the FC matrix to create a graph representation and calculate graph metrics^55,56^, we chose an edge density threshold of 5% for downstream analyses which ensures that the graph is sparse and free of negative correlations^57^. To demonstrate that our results are not dependent on the choice of threshold, we also showed results at a range of edge density similar to previous research^58^, an edge density of 1-5% at 1% intervals and 10%-90% at 10% intervals. This is achieved by finding the maximum spanning tree (MST) backbone first using the BCT toolbox function (backbone_wu.m) and continually adding edges with the largest correlation values until the desired edge density is reached^59^. This is to ensure graph connectedness at the sparsest densities.

Since strength and participation coefficient measures show a correlation with scan time, the remaining scan time after frame censoring was regressed out of these graph metrics to correct for the possible confound of individual differences in total scan time remaining after frame censoring (Supplementary Figure 3).

### Hub Disruption Index

To measure how the centrality of each region differs from a healthy reference, we chose NC match 1 to be the reference group. This choice is motivated by the fact that this group is the closest to what is usually considered young, healthy adults^22,23^, which has been used as a baseline for calculating hub disruption in prior studies^23^. The average nodal FC strength of the NC match 1 group was calculated and the percentage difference from this baseline average strength was calculated for each MC group (CDR = 0, CDR = 0.5, CDR≥1; Figure 3B) and the remaining two NC groups (NC match 2, NC match 3). While using a consistent baseline reference group enables comparison of the metric across groups^60^, we also ran a supplementary analysis using the age-matched NC groups as a reference for each MC group and obtained qualitatively the same result.

**Figure 3.**
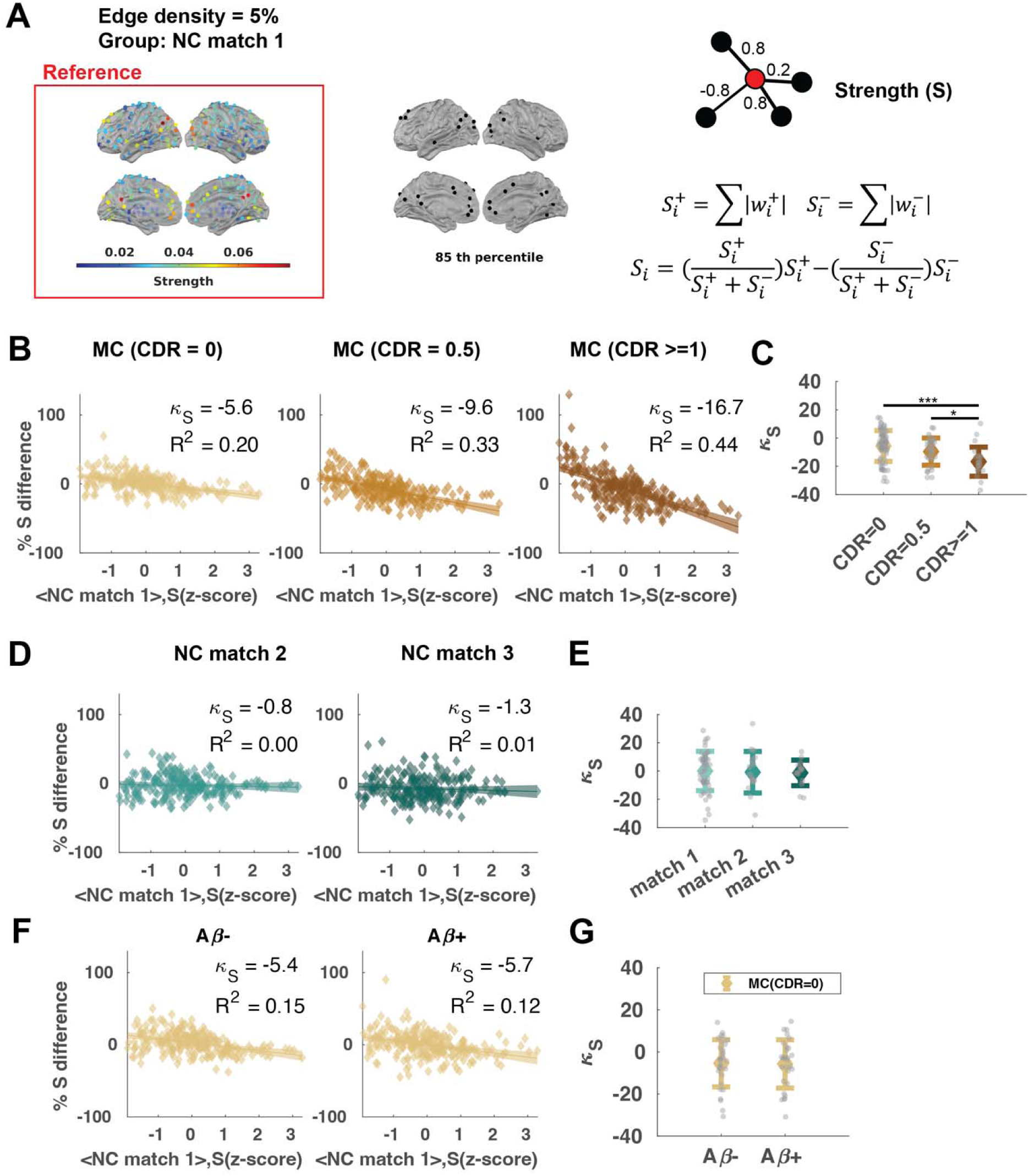
Hub disruption across CDR stages. **(A)** (Left) Distribution of average strength (S) across NC match 1 group, (Middle) nodes with S higher than the 85^th^ percentile. (Right) cartoon illustrating that strength is calculated by summing the weights across connected edges. **(B)** The % S difference against the baseline S Z-score in MC groups. **(C)** Individual hub disruption index (K_S_) for MC groups. **(D)** The % S difference against the baseline S Z-score in NC groups. E) Individual K_S_ for NC groups. **(F)** The % S difference against the baseline S Z-score in subsets of Aβ- and Aβ+ participants in the MC(CDR=0) group. **(G)** Individual K_S_ for Aβ- and Aβ+ participants in the MC(CDR=0) group. Shaded areas show 95% confidence interval. Error bars show mean and standard deviation.

We calculated a normalized hub disruption index adapted from prior work^60^ by fitting a linear regression slope (K_S_) where the dependent variable is the difference in strength for either the group average or an individual from the average strength of the reference group, which is then normalized by dividing the average strength of the reference group. The primary method by which selective hub disruption has been indexed in prior work is to calculate the slope of the linear regression model between the mean local network measures of a reference group, and the difference between that reference and the participant under study^23,60–63^(Figure 1A). A negative slope has been interpreted as the presence of hub disruption. In practice, this definition has difficulty distinguishing between a targeted attacks on network hubs vs. a random attack, as the correlation would appear negative under both models. An intuitive analogy is to examine the effect of a natural disaster on the affluent or impoverished areas. While affluent areas of a city lose more in absolute amounts, it is unclear whether they are *disproportionally more* affected, or it is merely an effect of starting from a higher baseline point. We, therefore, modified the “hub disruption index” to measure the *percentage* difference in connectivity strength versus baseline connectivity strength such that only targeted attacks on hubs would result in a negative linear regression slope (Figure 1B). We also standardized the baseline strength into Z-scores to increase the interpretability of the metric. In this way, we can unambiguously test the selective reduction in global connectivity at hubs and compare the hub disruption index quantitatively across disease stages. The mathematical equation is shown below:

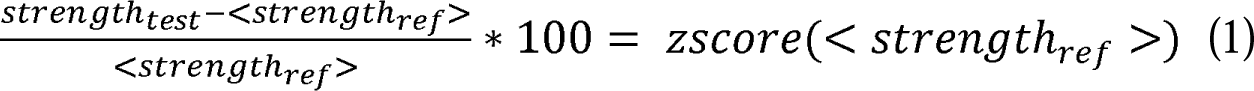

A more negative hub disruption index here indicates that the strengths in high-strength hubs are reduced by a larger proportion than in other low-strength regions, whereas a zero-hub disruption index indicates that the strength in high-strength regions and low-strength regions are changed to the same extent.

We used global connectivity strength as the primary measure of centrality due to its simplicity and strong correspondence to other measures in AD disease factors, e.g. hubs with high global FC have spatial correspondence with amyloid deposition^16,18,28^, tau burdens^30^ and metabolic factors^26,64^. However, there exist alternative definitions for hubs^20,57,65^. Specifically, some researchers argued that in functional networks, the participation coefficient, which captures the diversity of connections to different modules, is the key metric for regional importance or centrality^57^. On the other hand, within-module strength Z score is also an important measure of nodal centrality^54^. Therefore, we additionally examined hub disruption as defined by the participation coefficient and within-module Z-score using a similar asymmetric weighting^53^ with the BCT toolbox functions (participation_coef_sign.m) and custom MATLAB scripts, respectively.

### Cognitive Composite Score

We used a cognitive composite score (CCS)^66–68^, developed for use as an outcome measure in DIAN clinical trials, to measure the cognitive decline for each individual. CCS is a global summary of cognitive functions. Details of the calculation have been previously described^68^. Briefly, a cognitive composite score is calculated by averaging each test’s normalized scores by equal weight for 1) the DIAN Word List test delayed recall, 2) the delayed recall score from the Wechsler Memory Scale-Revised Logical Memory IIA subtest, 3) the Mini-Mental State Exam (MMSE), and 4) the Wechsler Adult Intelligence Scale-Revised Digit-Symbol Substitution test (Supplementary Table 4). Normalization was carried out with respect to the mean and SD reported in a population sample of 58 MCs with EYO<-15^68^. For analyses using the cognitive composite scores, we excluded one MC with a >1 year gap between psychometric tests and MRI sessions, and additionally 9 MC who did not complete all four tests.

### Positron emission tomography (PET) measures of cortical amyloid deposition

Amyloid-beta PET imaging with Pittsburgh Compound B (PiB) was performed using a bolus injection of [11C] PiB^40^. PET data were acquired using either a 70-min scan beginning at the start of the injection or a 30-min scan starting 40 min after the injection. Data were converted to regional standardized uptake value ratios (SUVRs) relative to the cerebellar grey matter using ROI generated in FreeSurfer^46^ with partial volume correction via a regional spread function . Amyloid positivity was defined as PiB partial volume corrected SUVR across the precuneus, prefrontal, gyrus rectus, and temporal FreeSurfer regions of interest (ROI)>1.42^69,70^.

### Statistical Models for Biomarkers

Generalized Additive Mixed Models were fit with the *gamm()* function from R package (*mgcv*) to examine the relationship between different biomarkers and the EYO. A smooth function was applied to the EYO separated by mutation carrier status (MC or NC), with sex, education as fixed effect covariates, and a random effect of family. The time of divergence between MC and NC was determined as the point where the predicted 95% simultaneous confidence interval starts to have no overlap^71^. For predicting the response of K_S_, we further include the residual motion measure (DVARS) as a covariate.

For the relationship between CCS and K_S_, we fit a linear mixed effects model with fixed effect covariates sex, education, DVARS and age and a random effect of family with the *<u>lmer()</u>* function from R package *lme4*.

### Statistical Tests and Visualization

All standard statistical tests (e.g., F-tests, t-tests, ANOVA) were performed with MATLAB R2020b or R (4.1.0). FDR^72^ was used for the correction of multiple comparisons at a significance level of 0.05.

Visualizations of regional graph theory metrics on the brain and FC matrix were generated using Network Level Analysis toolbox (Beta version) (https://github.com/mwheelock/Network-Level-Analysis), BrainNet Viewer toolbox^73^, and custom MATLAB and R scripts.

### Data availability

Data that support the findings of this study are available from DIAN at https://dian.wustl.edu/our-research/observational-study/dian-observational-study-investigator-resources/.

## Results

### MC and NC groups do not differ in demographic features and data quality

As designed, each of the MC and NC-matched CDR groups did not differ in age or EYO (Table 1). The matched groups also did not differ in DVARS or minutes of low-motion data. Not surprisingly, the CDR 0 groups did not differ on CCS, however, the MC and NC groups differed on CCS at CDR = 0.5 and CDR>=1.

### A selected subset of ROIs shows significant differences in strength from the healthy reference

We defined the average strength of ROIs in the young cognitively normal non-carrier group (NC match 1, N = 52) as a reference of hub centrality (Figure 3A). We first established that the number of participants and minutes of FC data were sufficient to obtain reliable group-level RSFC measures (Appendix A). Notably, we were able to identify the hubs described in literature^28,58,74^, e.g. precuneus/posterior cingulate, dorsolateral prefrontal cortex, supramarginal gyrus, medial prefrontal cortex (Figure 3A; Supplementary Figure 4). This is robust to the choice of edge density threshold ann d/or percentile cut-offs (Supplementary Figure 5). Additionally, we compared the ROI strengths between all MC groups and the reference using a two-sample t-test (FDR<0.05) (Supplementary Figure 6). Briefly, no ROI had significant difference in strength between MC (CDR=0) and the reference. In MC (CDR=0.5), 19 ROIs covering the superior frontal gyrus, precuneus, middle temporal gyrus, middle occipital gyrus, middle frontal gyrus, inferior parietal lobule, inferior occipital gyrus, fusiform gyrus and cuneus have significantly lower strength compared to the reference. In MC (CDR≥1), three ROIs (in the insula, thalamus and parahippocampal gyrus) showed significant higher strength compared to the reference, and 30 ROIs (in angular gyrus, anterior cingulate, claustrum, cuneus, fusiform gyrus, inferior parietal lobule, inferior temporal gyrus, insula, medial frontal gyrus, middle occipital gyrus, middle temporal gyrus, parahippocampal gyrus, postcentral gyrus, posterior cingulate, precuneus, superior frontal gyrus, superior temporal gyrus and thalamus) showed significantly lower strength compared to the reference. On the other hand, none of the ROIs in NC match 2 or NC match 3 groups shown significant differences in strength from NC match 1 (Supplementary Figure 7).

### Hub disruption increases with CDR stage, not age

We measured the group-level hub disruption index by calculating the percentage difference from the reference for the mean strength in each of the MC groups (CDR=0, CDR=0.5, and CDR ≥ 1) (Figure 3B; Supplementary Figure 8). The group-level hub disruption index for all three MC CDR groups was significantly different from zero (Table 2). In addition, K_S_ became increasingly more negative across CDR stages. The hub disruption index (a.k.a. regression slope in Figure 3B) is significantly different between MC (CDR = 0.5) and MC (CDR = 0) (*F*(1,488) = 12.0, *P*<0.001, partial 77^2^ = 0.024), between MC (CDR=0.5) and MC (CDR≥1) (*F*(1,488) = 22.6, *P*<0.001, partial 77^2^ = 0.044), and between MC (CDR=0) and MC (CDR≥1) (*F*(1,488) = 61.6, *P*<0.001, partial 77^2^ = 0.112). Our results are qualitatively replicated at a wide range of threshold choices (Supplementary Figure 9). On average, nodes in the cingulo-opercular network showed the highest baseline strength and largest % strength difference from baseline across multiple thresholds (Supplementary Figure 10). In addition to the group-level hub disruption index, we calculated the hub disruption index for each participant in the MC and NC groups. All MC groups had a hub disruption index that differed from zero (FDR-adjusted *P*<0.001) while no NC group had a hub disruption index that differed from zero (FDR-adjusted *P*>0.05) (Supplementary Figure 11, Supplementary Table. 5). Specifically, for MC, a one-way ANOVA demonstrated that the hub disruption index differed across the CDR groups (*F*(2,118) =8.8, *P*<0.001, 77^2^ = 0.130). Post-hoc two-sample t-tests with FDR correction revealed significant group differences (*t*(87) = 4.03, *P* = 0.002, Cohen’d = 1.02) between CDR=0 (M = -5.6, SD = 11.1) and CDR≥1 participants (M=-16.7, SD=10.3), and between CDR=0.5 (M = -9.6, SD = 9.6) and CDR≥1 participants (*t*(50) = 2.52, *P*=0.03, Cohen’s d = 0.72) (Figure 3C).

**Table 2.** Group-level hub disruption (using metrics in NC match1 as baseline) across CDR stages in MC and across age in NC (FDR-adjusted).

| Strength (S) |  |  |  |  |  |
| --- | --- | --- | --- | --- | --- |
| | Group | $\kappa_S$ | F (1,244) | p | $R^2$ |
| MC | CDR = 0 | -5.6 | 59.1 | <0.001 | 0.20 |
|  | CDR = 0.5 | -9.6 | 118.1 | <0.001 | 0.33 |
|  | CDR ≥ 1 | -16.7 | 192.2 | <0.001 | 0.44 |
| NC | match 2 | -0.8 | 0.78 | 0.38 | 0.003 |
|  | match 3 | -1.3 | 1.46 | 0.28 | 0.006 |
| Participation Coefficient (Pc) |  |  |  |  |  |
| | Group | $\kappa_S$ | F (1,244) | p | $R^2$ |
| MC | CDR = 0 | 3.3 | 17.8 | <0.001 | 0.07 |
|  | CDR = 0.5 | 1.7 | 2.5 | 0.19 | 0.01 |
|  | CDR ≥ 1 | -1.6 | 1.0 | 0.34 | 0.44 |
| NC | match 2 | 2.1 | 4.7 | 0.08 | 0.02 |
|  | match 3 | -1.0 | 0.9 | 0.34 | 0.003 |
| Within-module Strength Z-score (Z) |  |  |  |  |  |
| | Group | $\kappa_S$ | F (1,244) | p | $R^2$ |
| MC | CDR = 0 | -5.0 | 12.6 | <0.001 | 0.05 |
|  | CDR = 0.5 | -7.2 | 15.5 | <0.001 | 0.06 |
|  | CDR ≥ 1 | -12.7 | 22.1 | <0.001 | 0.08 |
| NC | match 2 | -2.1 | 1.6 | 0.21 | 0.01 |
|  | match 3 | -3.3 | 2.9 | 0.11 | 0.01 |

Next, we asked whether this observation can be explained by increasing age. We calculated the hub disruption index for the age-matched NC groups 2 and 3 with the same procedure (Figure 3D). The group-level hub disruption index for NC groups did not significantly differ from zero (Table 2). At the individual level, there were no differences among NC groups (one-way ANOVA, *F*(2,81) = 0.07, *P* = 0.93) (Figure 3E), nor a *2* significant relationship between K_S_ and age in NC (linear regression, f3 = 0.01, *R* <0.001, *F*(2, 82) = 0.0072, *P* = 0.933).

Changes in AD biomarkers often precedes dementia symptoms in AD^38,75,76^. A subset of the MC (CDR=0) group can be classified as amyloid beta positive (Aβ+) (*N* = 29/60) according to their amyloid PET results (Methods). With one-sample t-tests with FDR correction, we found that both groups had hub disruption index significantly lower than 0 (Aβ-: M = -5.4, SD = 11.2, Cohen’s d = -0.48, *t*(30) = -2.7, *P* = 0.012; Aβ+: M = -5.7, SD = 11.5, Cohen’s d = -0.50, *t*(28) = -2.6, *P* = 0.012). However, there was no significant difference in hub disruption index between the Aβ- and Aβ+ groups (two-sample t-test, two-tailed *P* = 0.92) (Figure 3F-G). Changes in amyloid-beta accumulation in PET imaging often precedes dementia symptoms in AD^38,75,76^. A subset of the MC (CDR=0) group can be classified as amyloid beta positive (Aβ+) (*N* = 29) according to their amyloid PET results (Methods). With one-sample t-tests with FDR correction, we found that both groups have hub disruption index significantly lower than 0 (Aβ-: M = -5.4, SD = 11.2, Cohen’s d = -0.48, *t*(30) = -2.7, *P* = 0.012; Aβ+: M = -5.7, SD = 11.5, Cohen’s d = -0.50, *t*(28) = -2.6, *P* = 0.012). However, there was no significant difference in hub disruption index between the Aβ- and Aβ+ groups (two-sample t-test, two-tailed *P* = 0.92) (Figure 3F-G).

### Hub disruption is best explained by differences in regional global connectivity

To understand the key drivers of hub vulnerability in ADAD, we calculated the hub disruption index using two alternative measures based on their network membership instead of global connectivity strength: 1) the within-module connectivity rank (within-module strength Z-score, *Z*) and 2) the connectivity diversity (Participation Coefficient, *Pc*) (Figure 4A). Overall, both participation coefficient and within module Z-score effects were less sensitive to ADAD progression than using the global connectivity strength as the reference. Thus, we focused subsequent analyses on the hub disruption index with regards to the global connectivity strength. Detailed statistics can be found in Table 2 and Appendix B.

**Figure 4.**
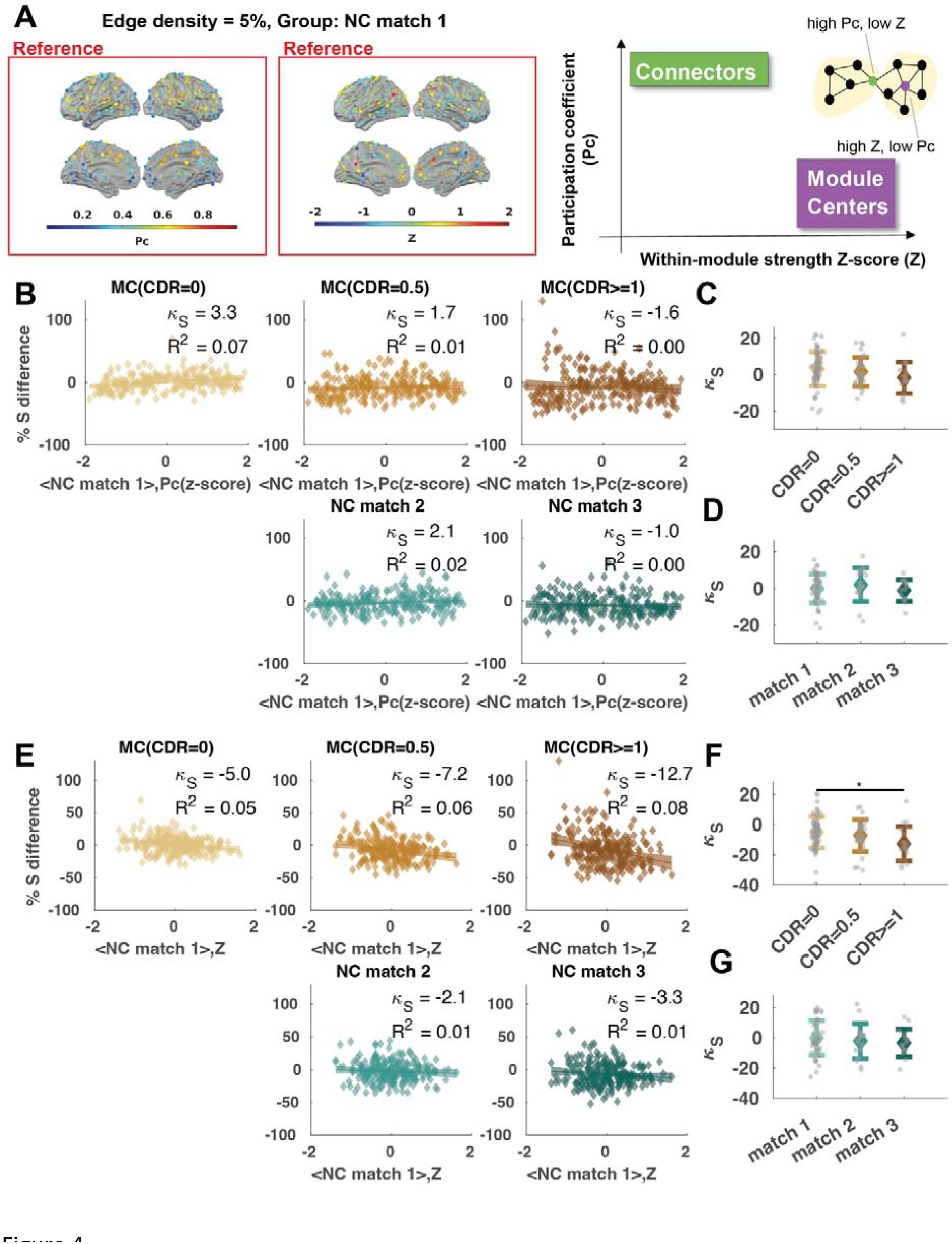
Hub disruption across CDR stages at module centers versus connectors. **(A)** (Left) Distribution of average participation coefficient (Pc) across NC match 1 group, (Middle) distribution of within strength Z-score (Z) across NC match 1 group, (Right) cartoon illustrating the representation of module centers and connectors on a graph. Module centers are nodes with high Z and connectors are nodes with high Pc. **(B)** The % S difference against the baseline Pc Z-score for hub disruption calculation. **(C)** Individual hub disruption index (K_S_) for MC with respect to the group average Pc Z-score at NC match 1. **(D)** Individual hub disruption index (K_S_) for NC with respect to the group average Pc Z-score at NC match 1. **(E)** The % S difference against the baseline Z for hub disruption calculation. **(F)** Individual hub disruption index (K_S_) for MC with respect to the group average Z at NC match 1. G) Individual hub disruption index (K_S_) for NC with respect to the group average Z at NC match 1. Lines show linear fit and shaded areas indicate the 95% CI.

### Hub disruption index diverges between mutation carriers and non-carriers at an earlier EYO than the separation of general cognitive performance

Generalized Additive Mixed Models were fit to examine the relationship between hub disruption or other biomarkers and the EYO, as well as to obtain the point of divergence between MC and NC. For the hub disruption index (K_S_), this was calculated to be EYO = -7.9 years (Figure 5A). In comparison, the total cortical amyloid deposition measured as PiB SUVR ratio diverged at EYO = -14.8 (Figure 5B), and the cognitive composite score measure diverged at EYO = -6.8 years (Figure 5C). Thus, we found that the divergence of hub disruption index preceded the divergence of cognitive performance measure and followed the earlier stage of amyloid deposition.

**Figure 5.**
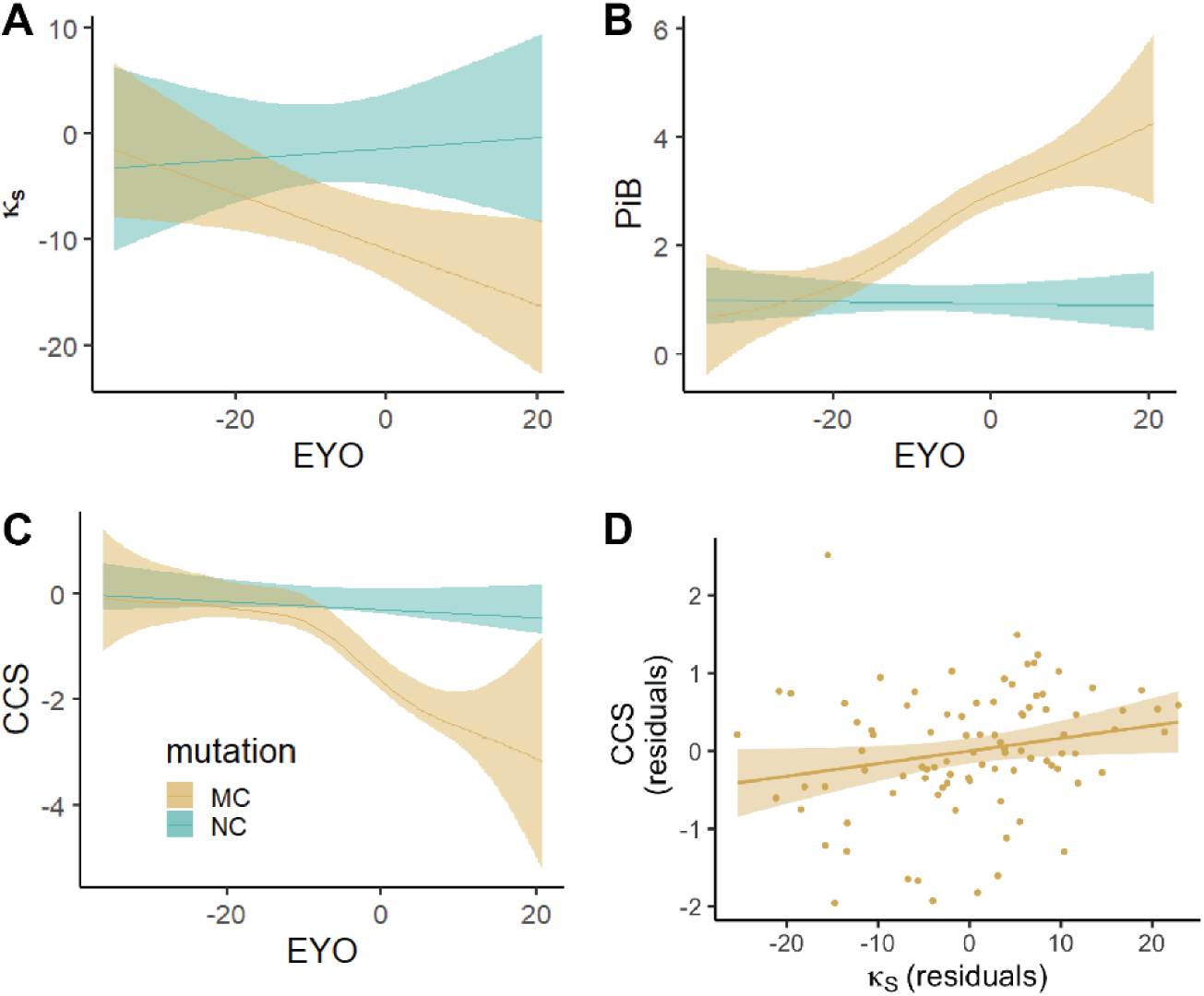
Change in Biomarkers Across Estimated Years to Symptom Onset (EYO) for mutation carriers (MC) and non-carriers (NC). **(A)** The hub disruption index in strength (K_S_) against EYO, **(B)** The total cortical amyloid deposition measured with PiB against EYO, **(C)** The Cognitive Composite Score (CCS) against EYO. **(D)** The CCS against K_S_ after regressing out potential confounding variables from both. The line and shaded areas show the predicted response values and the confidence intervals for the fitted responses from a generalized additive model at 95% interval calculated at each observation. For privacy reasons, the extreme EYO values (EYO<−20 and >10) were not displayed but were used in model-fitting.

### Greater hub disruption is correlated with worse general cognition

Lastly, we found there existed a positive correlation between K_S_ and CCS (*r*=0.3, *t*(110)=3.27, *P*=0.001). We further examined whether an individual’s hub disruption could explain unique variance in the cognitive composite score of individual MCs after controlling for potentially confounding covariates (age, sex, years of education, motion in scan measured by DVARS; and family as a random effect. The hub disruption index was positively related with cognitive composite scores at the edge threshold of 5% (f3_K_S_=0.02±0.01, *t*(105)=2.53, *P*=0.013) and across different edge thresholds (Table 3, Figure 5D), suggesting that greater hub disruption (a.k.a. more negative hub disruption index) correlated with worse general cognition.

**Table 3.** Regression of hub disruption index on Cognitive Composite Score (CCS). Response: Cognitive Composite Scores (CCS). β, coefficient of regression. ***p<0.001, **p<0.01, *p<0.05. Random effect: family.

| Edge threshold | $\beta_{\_KS}$ | $\beta_{\_Education}$ | $\beta_{\_Age}$ | $\beta_{\_Sex \text{ (Male)}}$ | $\beta_{\_DVARs}$ |
| --- | --- | --- | --- | --- | --- |
| 5% | 0.02* | 0.11*** | -0.06*** | -0.18 | -0.04 |
| 10% | 0.03** | 0.11*** | -0.06*** | -0.18 | -0.05 |
| 20% | 0.05** | 0.11*** | -0.06*** | -0.19 | -0.06 |
| 30% | 0.07** | 0.11*** | -0.06*** | -0.20 | -0.07 |
| 40% | 0.09** | 0.10*** | -0.06*** | -0.21 | -0.07 |

## Discussion

Consistent with a targeted attack on hubs model, the proportion of reduction in FC at individual regions in ADAD was positively related to the total global connectivity of that region in the unaffected family members of ADAD participants. This preferential disruption of hub connectivity increased with CDR stage but not age, is best explained by global connectivity, less so to the within-module connectivity rank, and not to the diversity of connectivity across resting-state networks. This preferential disruption of hub connectivity is seen at all stages of disease progression in ADAD MC and starts to differentiate MC and NC at about EYO = 8 years. Additionally, greater (more negative) hub disruption was associated with worse general cognition after controlling for relevant covariates.

### Progressive hub disruption is consistent with popular network failure models of AD

Prior studies endorse a cascading network failure starting from the posterior default mode network (DMN) and progressing to the anterior and ventral DMN^13^. Our results complement this observation by providing a possible underlying mechanism for this cascading process and extending it beyond default mode network, whereby the vulnerability of regions to the reduction in FC is dependent on their centrality in the whole brain network. Nodes that have the highest centrality (e.g., posterior default mode network, Figure 3A) were among the first to show decreased FC, consistent with existing literature^13^. Over time, changes in processing burden shifts from one hub to other hubs, further enhancing the aberrant amyloid-f3 precursor protein processing and amyloidosis in the other hubs^13^, consistent with our progressive increase in hub disruption with disease. Admittedly, FC is an indirect measure of connectivity across regions, and future research could investigate hub vulnerability in structural networks using diffusion MRI.

### Hub disruption is best explained by global connectivity

Despite the wide use of “functional hubs” in the literature, what defines a functional hub has not reached a consensus^20,57,65^. Hubs can be described in terms of their network membership (e.g., default mode network), where connectors are important for communication between networks and module centers are important for communication within networks. The nodes with high abundance of intermodule connections (connectors) form a structural rich club^80^, which are also known to be affected in AD^81^, although this concept is not universally accepted (other studies suggest that the highly rich-club core was preserved and the disruptions started in the periphery^82^). Previous literature on brain lesion patients suggested that the integrity of brain network organization is severely compromised when damage is in connectors but not module centers^83^. Other studies also report differential outcomes in network structure when damage is localized to module centers or connectors^34^. One recent study has also suggested that the amyloid-beta accumulation rate was faster at connectors^19^. On the other hand, our results here suggested that hub disruption is best explained by differences in global connectivity across regions, rather than their roles to communicate between or within networks. This is in line with the hypothesis that high metabolic demands associated with high global connectivity may trigger downstream cellular and molecular events that result in neurodegeneration^13^, conveying preferential/selective vulnerability. For example, calcium instability caused by amyloid beta peptides may render human cortical neurons vulnerable to excitotoxicity^84^, and this could result in further neurodegeneration in AD^85^.

### Hub disruption predates cognitive changes but follows amyloid PET changes

The effectiveness of a biomarker can be evaluated based on its ability to detect early indications of pathology prior to disease onset. Investigating the initial stages of decline in healthy brains compared to those AD offers substantial potential for early identification before AD symptoms manifest. Because of the highly consistent familial disease onset for ADAD, we were able to compare this biomarker across EYO and other disease-related changes including cognitive composite scores and cortical amyloid deposition. We found that the hub disruption index first demonstrated a divergence between groups ∼8 years before EYO—much earlier than the divergence in global FC signature (∼4 years)—between converters and non-converters in sporadic AD^71^. Hub disruption index also diverges between MCs and NCs before the changes in the general cognitive score (∼7 years), but after hypo-metabolism (∼10 year), and increased concentrations of cerebral spinal fluid tau protein (∼15 year)^38^, and the amyloid PET changes (∼15 year). This is consistent with previously hypothesized disease progression where the FC disruption follows from amyloid-beta deposition and potentially excessive chronic activity^24,86^, and eventually contributes to cognitive impairment (Supplementary Figure 16). We did not find significant differences between the hub disruption index in Aβ+ and Aβ-participants in the MC (CDR=0) group, despite Aβ+ participants having slightly negative hub disruption index. However, this lack of a difference between Aβ- and Aβ+ individuals should be viewed cautiously given the modest sample sizes. We do note the limitation that our EYO calculation is based on mutation and parental symptom onset and may not precisely reflect the true EYO. Therefore, the best practice is to interpret the EYO years in relative terms for different biomarkers instead of taking it at purely its face value. We did not find significant differences between the hub disruption index in Aβ+ and Aβ-participants in the MC (CDR=0) group, despite Aβ+ participants having slightly negative hub disruption index. However, this lack of a difference between Aβ- and Aβ+ individuals should be viewed cautiously given the small sample size for the comparison.

### Comparison to other network topology studies in AD

Other studies of network topology in AD have examined global graph theory measures such as small-worldness, global clustering coefficient, and characteristic path length^58,87^. However, those measures are generally sensitive to network sparsity and require a careful choice of null models^88^. Further, it is hard to interpret the biological relevance for those global measures. In contrast, hubs with high global FC have been linked to amyloid deposition^16,18,28^, tau burden^30^ and metabolic factors^26,64^. They also overlap with the regions that demonstrated high heritability^89^. Therefore, our research on hub vulnerability is literature-driven with an attempt to link abstract network topology measures to molecular and cellular pathologies.

### Implications for AD research, prevention, and treatment

We found that hub disruption, or increased vulnerability to reduced FC at highly central hub regions, is prevalent across the course of ADAD, with increasing severity as the disease progresses. Our results here have key implications for future AD research and therapeutics development: we provided a testable hypothesis where targeted pharmacological manipulation, non-invasive stimulation, or behavioral training to alter neuronal excitability^90^ especially at hub regions could alter the progression of AD. Existing research has demonstrated in an awake rodent model that acute inactivation of a hub region (dorsal anterior cingulate cortex) has profound effects on the whole network^91^. Future studies in animal models of AD could further validate this with optogenetic and chemogenetic manipulations. Furthermore, previous literature has suggested that “restoration of the topology of resting-state FC may aid in cognitive repair and recovery”^32,92^, and those can be further tested in future studies.

On the other hand, we found that hub disruption is positively related to the cognitive composite scores after considering the effect of age, sex, years of education, and average motion of retained frame. And the separation of hub disruption between MC and NC starts shortly after the increased levels of cortical amyloid deposition and at around the same time as preclinical measures of cognitive decline. This indicates that our new measure of resting-state FC change has the potential to act as a non-invasive, low-cost, and accessible biomarker especially given compared to CSF and PET for prevention studies and clinical trials to aid the development of new treatments and monitor their effectiveness. Other biomarkers focusing on DMN network failure have been proposed^93^, but our measure is conceptually straightforward, easy to calculate, and biologically intuitive. In addition, previous measures have focused on distinguishing AD patients from controls, whereas the current study mapped a progressive relationship between FC and centrality across the clinical dementia stages.

### Limitations and future directions

While we concluded that increasing hub disruption was related to disease progression and not aging, participants involved in this study were relatively young (18-69 years). It is still possible that a similar increase in hub vulnerability would be observed at a much older age, as seen in other age-related changes in FC^94,95^. Notably, another study using cognitively normal adults from OASIS-3 (42-95 years) seemed to show the opposite result to the current study^19^, whereby functional hubs were particularly vulnerable to the higher annual accumulation of amyloid beta but have a slower FC decrease than non-hub regions. However, there are also several important methodological differences between that study and ours: 1) they employed the GLASSO algorithm to estimate FC with only direct connections while we used the simple Pearson’s correlation, and 2) they define hubs as regions with high participation coefficients and we found that at certain edge density threshold, the strength and the participation coefficient of a node could be negatively correlated. Additionally, even though previous work has found comparable FC changes in ADAD to sporadic AD (Smith et al., 2021; Strain et al., 2022; Wheelock et al., 2023), our results are yet to be confirmed in sporadic AD. Further validations on longitudinal changes, and on subjects with more imaging data are needed to assess whether hub disruption could be a reliable biomarker of individual disease progression in AD. Furthermore, future investigations in large brain-wide single-cell transcriptome data (e.g. Allen Human Brain Atlas) may be useful in linking the hub vulnerability to the underlying biological mechanisms^96,97^.

## Conclusions

We investigated the relationship between FC differences across ROIs and baseline centrality measures. We demonstrated that hubs with high global connectivity are especially vulnerable to reduction in FC in individuals with ADAD, consistent with a targeted attack on hubs model. Moreover, this disruption of hub connectivity becomes more severe with increasing CDR stage and occurs around 8 years before symptom onset, slightly preceding cognitive changes but following amyloid PET changes, indicating the early and progressive nature of hub vulnerability in AD. Interestingly, our results also suggest that the preferential disruption of FC in hub regions is more related to global connectivity rather than within-module connectivity or diversity of connectivity across networks. These findings provide insights into the complex dynamics of brain network dysfunction in AD and the critical role of hubs in this process.

## Acknowledgements

Data collection and sharing for this project was supported by The Dominantly Inherited Alzheimer Network (DIAN, U19AG032438) funded by the National Institute on Aging (NIA), the Alzheimer’s Association (SG-20-690363-DIAN), the German Center for Neurodegenerative Diseases (DZNE), Raul Carrea Institute for Neurological Research (FLENI), Partial support by the Research and Development Grants for Dementia from Japan Agency for Medical Research and Development, AMED, and the Korea Health Technology R&D Project through the Korea Health Industry Development Institute (KHIDI), Spanish Institute of Health Carlos III (ISCIII), Canadian Institutes of Health Research (CIHR), Canadian Consortium of Neurodegeneration and Aging, Brain Canada Foundation, and Fonds de Recherche du Québec – Santé. This manuscript has been reviewed by DIAN Study investigators for scientific content and consistency of data interpretation with previous DIAN Study publications. We acknowledge the altruism of the participants and their families and contributions of the DIAN research and support staff at each of the participating sites for their contributions to this study.

During the preparation of this work the author(s) used ChatGPT in order to increase the clarity and conciseness of the language. After using this tool/service, the author reviewed and edited the content as needed and take full responsibility for the content of the publication.

We would like to additionally thank Aaron Tanenbaum for processing the MRI data, Dr. Julie Wisch and Dr. Tyler Blazey and Dr. Matt Welhaf for their suggestions and assistance in estimating the divergence of metrics between mutation carriers and non-carriers, as well as Dr. Nilanjan Chakraborty, Sayan Das, Dr. Aishwarya Rajesh, and Jiaqi Li for their discussion on the mathematical rigor.

## Supplementary material

Supplementary material is available at *Brain* online

## Appendix 1 Dominantly Inherited Alzheimer Network

Randall Bateman, Alisha J. Daniels, Laura Courtney, Eric McDade, Jorge J. Llibre-Guerra, Charlene Supnet-Bell, Chengie Xiong, Xiong Xu, Ruijin Lu, Guoqiao Wang, Yan Li, Emily Gremminger, Richard J. Perrin, Erin Franklin, Laura Ibanez, Gina Jerome, Elizabeth Herries, Jennifer Stauber, Bryce Baker, Matthew Minton, Carlos Cruchaga, Alison M. Goate, Alan E. Renton, Danielle M. Picarello, Tammie Benzinger, Brian A. Gordon, Russall Hornbeck, Jason Hassenstab, Jennifer Smith, Sarah Stout, Andrew J. Aschenbrenner, Celeste M. Karch, Jacob Marsh, John C. Morris, David M. Holtzman, Nicolas Barthelemy, Jinbin Xu, James M. Noble, Sarah B. Berman, Snezana Ikonomovic, Neelesh K. Nadkarni, Gregory S. Day, Neill R. Graff-Radford, Martin Farlow, Jasmeer P. Chhatwal, Takeshi Ikeuchi, Kensaku Kasuga, Yoshiki Niimi, Edward D. Huey, Stephen Salloway, Peter R. Schofield, William S. Brooks, Jacob A. Bechara, Ralph Martins, Nick C. Fox, David M. Cash, Natalie S. Ryan, Mathias Jucker, Christoph Laske, Anna Hofmann, Elke Kuder-Buletta, Susanne Graber-Sultan, Ulrike Obermueller, Johannes Levin, Yvonne Roedenbeck, Jonathan V[glein, Jae-Hong Lee, Jee Hoon Roh, Raquel Sanchez-Valle, Pedro Rosa-Neto, Ricardo F. Allegri, Patricio Chrem Mendez, Ezequiel Surace, Silvia Vazquez, Francisco Lopera, Yudy Milena Leon, Laura Ramirez, David Aguillon, Allan I. Levey, Erik C.B Johnson, Nicholas T. Seyfried, John Ringman, Anne M. Fagan, and Hiroshi Mori.

