## Supplementary Materials for "Increasing hub disruption parallels dementia severity in autosomal dominant Alzheimer disease"

### Supplementary Table

**Supplementary Table 1.** Sample Characteristics (mutation carriers and non-carriers)

| Measure | Non-Carrier (N=85) | Mutation Carrier (N=122) | Test statistic | df | p-value |
|---|---|---|---|---|---|
| | | | Chi-Square | | |
| Sex (M/F) | 34/51 | 58/64 | 3.70 | 1 | 0.28 |
| CDR (0, 0.5, ≥1) n (%) | 85 (100%), 0 (%), 0 (%) | 70 (57.38%), 32 (26.23%), 20 (16.39%) | 48.38 | 2 | <0.001 |
| | | | Mann-Whitney U | | |
| | median | median | | z | p-value |
| Age (yrs) | 37.9 | 40.2 | 8782 | 0.1368 | 0.8912 |
| Education (yrs) | 15 | 14 | 9643 | 1.9109 | 0.056 |
| EYO | -8.9 | -6.7 | 8504 | 0.7913 | 0.4287 |
| CCS[a] | 0.06 | -0.81 | 1065 | 5.5011 | <0.001 |

a Missing 10 participant.

Medians and Mann Whitney test statistic reported (Shapiro-Wilk test of normality p<0.001)
EYO; estimated years from expected symptom onset; CCS, Cognitive Composite Score; CDR, Clinical Dementia Rating.

**Supplementary Table 2.** Sequence details

| | |
|---|---|
| MP-RAGE | TR =2300ms; TE = 2.95ms; flip angle 9˚; TI= 900ms; 256x240 acquisition matrix; 176 sagittal slices 1x1x1.2mm resolution. |
| fMRI | TR=2200ms; TE=27ms; flip angle=90˚;64x64 acquisition matrix, 36 axial slices in ascending interleaved order; 4mm isotropic resolution. |
| | TR=2200ms; TE=30ms; flip angle=80˚; 64x58 acquisition matrix; 36 axial slices in ascending interleaved order; 3.3mm isotropic resolution. |
| | TR=3000ms; TE=30ms; flip angle=80˚; 64x58 acquisition matrix; 48 axial slices in ascending interleaved order; 3.3mm isotropic resolution. |

**Supplementary Table 4**. Tests for assessing general cognitive functions to give cognitive composite scores(Wang et al. 2018)

| Test | Explained | Range |
|---|---|---|
| MINI MENTAL STATE EXAM(Folstein, Folstein, and McHugh 1975) | Scored according to the UDS guidebook. | 0-30 |
| WMS-R LOGICAL MEMORY IIA - DELAYED(Wechsler 1987) | Administered after WAIS-R Digit Symbol in prescribed UDS order, and scored according to WMS-R manual | 0-25 |
| WAIS-R DIGIT SYMBOL(Wechsler 1981) | This is an enlarged Digit Symbol form that measures 15 x 24 cm rather than 9.5 x 13 cm as in the standard WAIS-R. Otherwise administered and raw scored according to WAIS-R manual. | 0-93 |
| WORD LIST RECALL - Delayed | Number of words from word list recalled after delay interval. | 0-16 |
| Source: DIAN CODEBOOK V1.5 1_1_2019_PDF | | |

**Supplementary Table 5.** Comparing the distribution of individual hub disruption index to zero

| **Groups** | Mean | S.D. | Cohen's d | d.f. | t | p |
|---|---|---|---|---|---|---|
| MC (CDR=0) | -5.6 | 10.9 | -0.51 | 68 | -4.28 | <0.001 |
| MC (CDR=0.5) | -9.6 | 9.6 | -1.00 | 31 | -5.64 | <0.001 |
| MC (CDR>=1) | -16.7 | 10.3 | -1.62 | 19 | -7.25 | <0.001 |
| NC (match 1) | 0 | 13.9 | 0 | 51 | 0 | 1.0 |
| NC (match 2) | -0.8 | 14.7 | -0.06 | 16 | -0.24 | 1.0 |
| NC (match 3) | -1.3 | 9.1 | -0.15 | 14 | -0.57 | 1.0 |

### Supplementary Figures

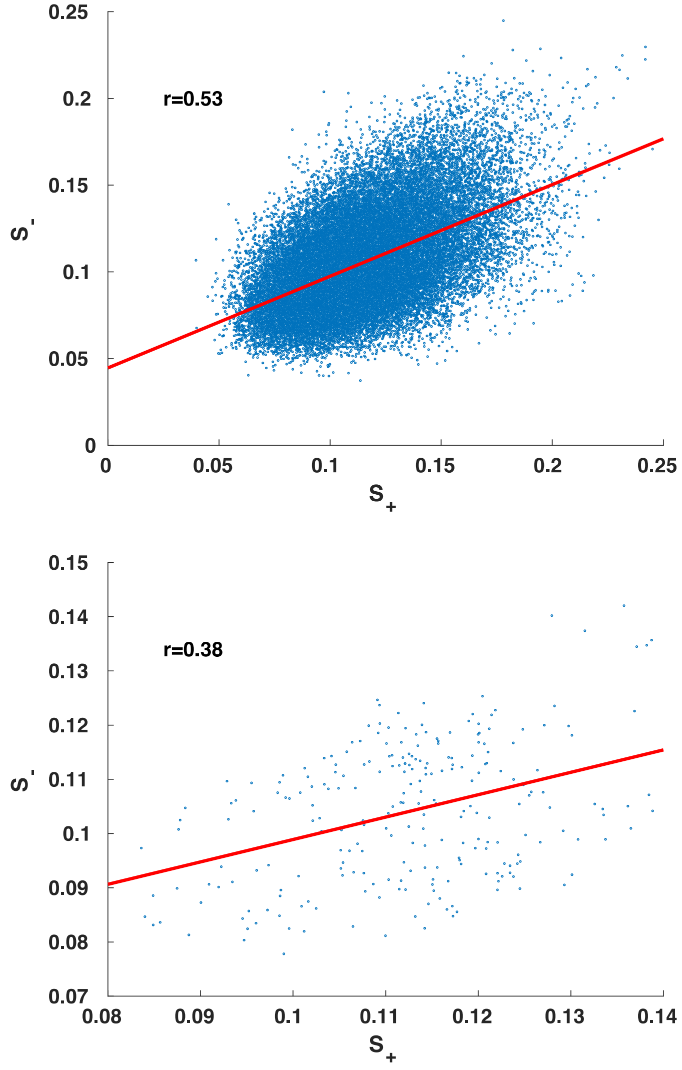


**Supplementary Figure 1. Nodal strength (S) calculated with positive and negative connections in the full matrix without threshold are highly correlated.** (Top) Across all subjects. Pearson's correlation, r= 0.53. (Bottom) Group average. Pearson's correlation, r = 0.38. $S_+$ and $S_-$ are the average of positive weights and negative weights, respectively. Red line shows linear fit.

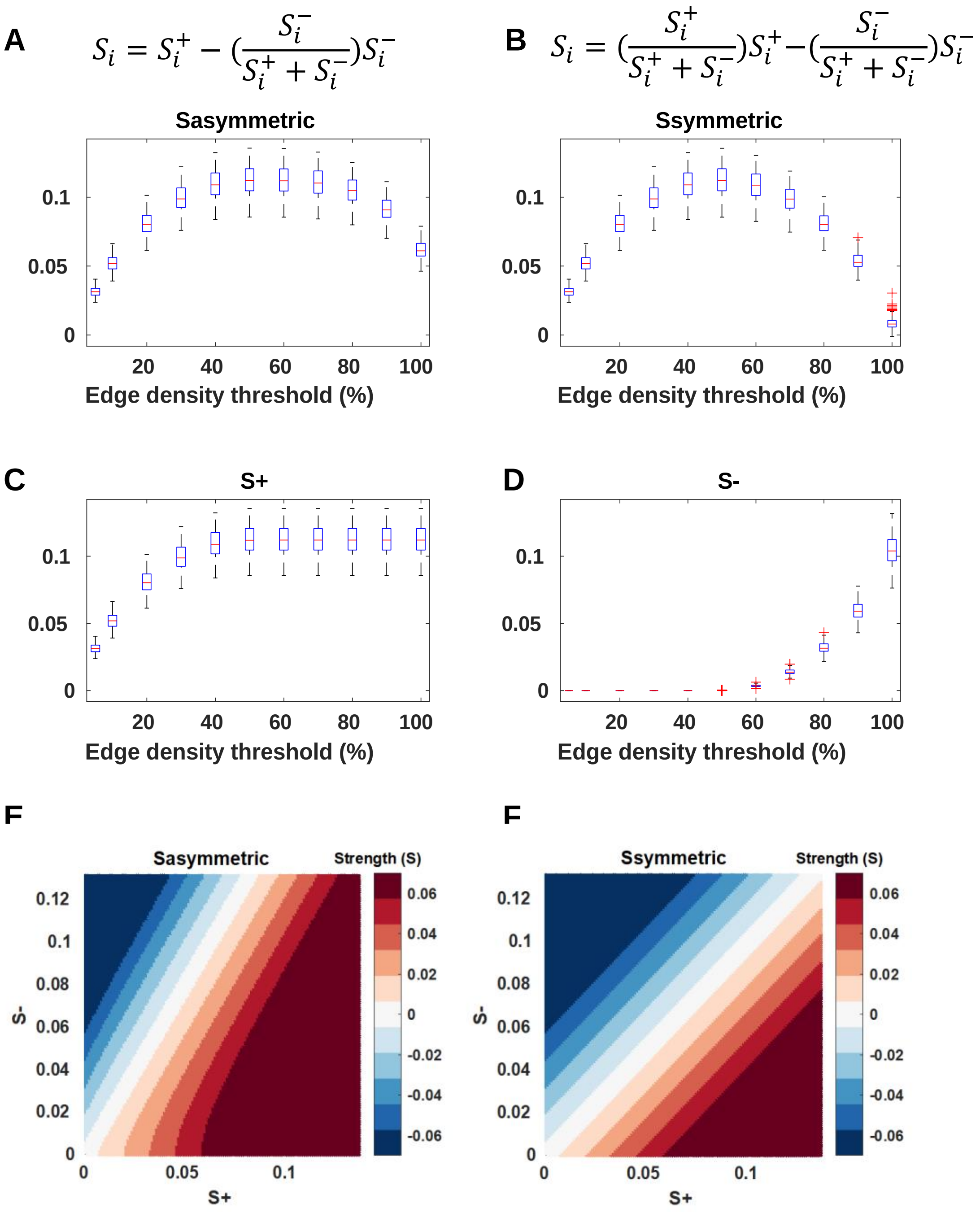


**Supplementary Figure 2. The calculation of nodal strength from positive and negative strengths.** a) Formula for asymmetric strength calculation and the asymmetric strength size at different densities. b) Formula for symmetric strength calculation and the symmetric strength size at different densities. c) Strength for positive edges at different densities. d) Strength for negative edges at different densities. e) Asymmetric strength for combinations of S+ and S-. f) Symmetric strength for combinations of S+ and S-.

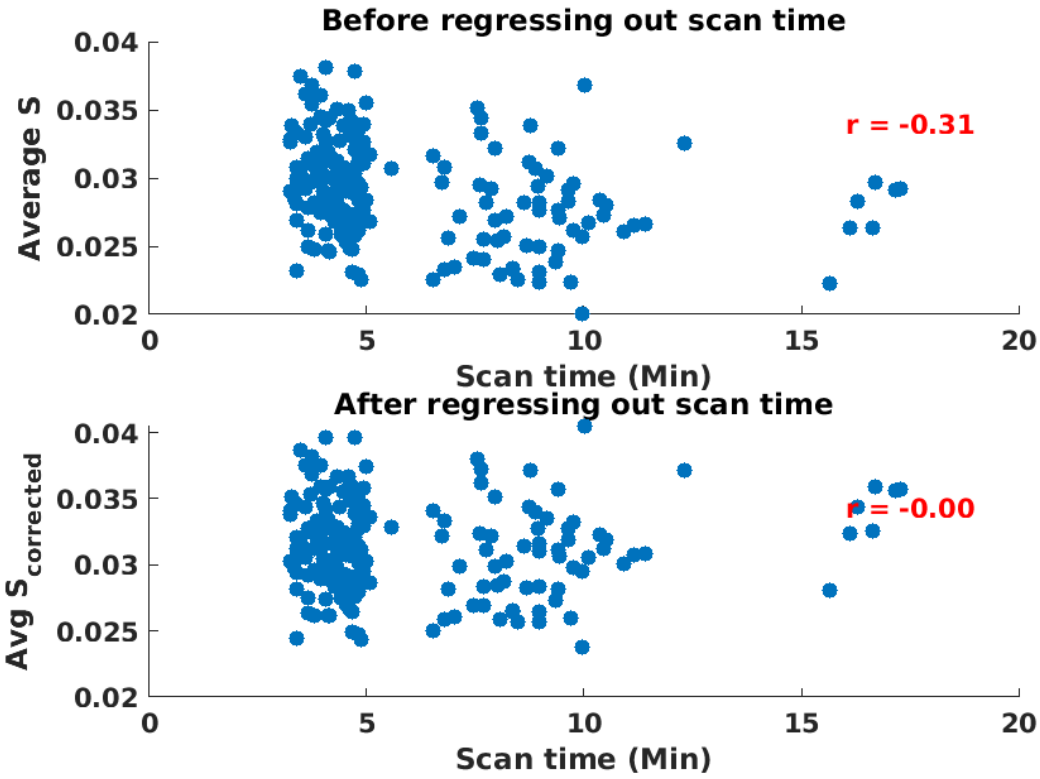


**Supplementary Figure 3. Correlation between Average nodal strength (S) and scan time.** (Top) before and (Bottom) after linear regression to correct for the correlation between average nodal strength and scan time. Edge density = 5%.

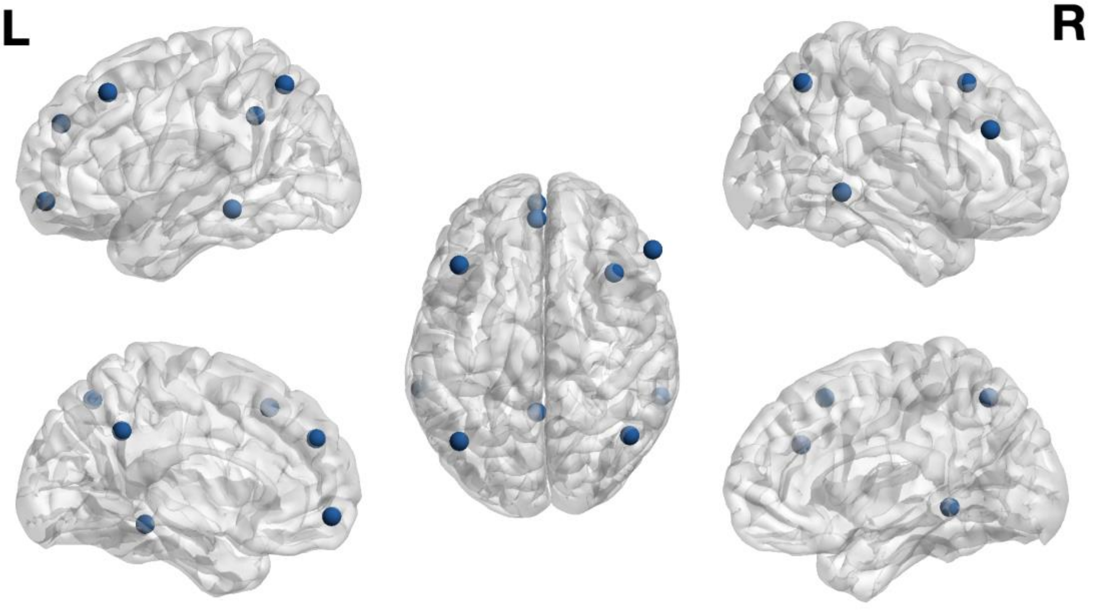


**Supplementary Figure. 4 Hubs identified in Buckner et al., 2009 J.Neurosci.** Nodes generated with the MNI coordinates from the original paper.

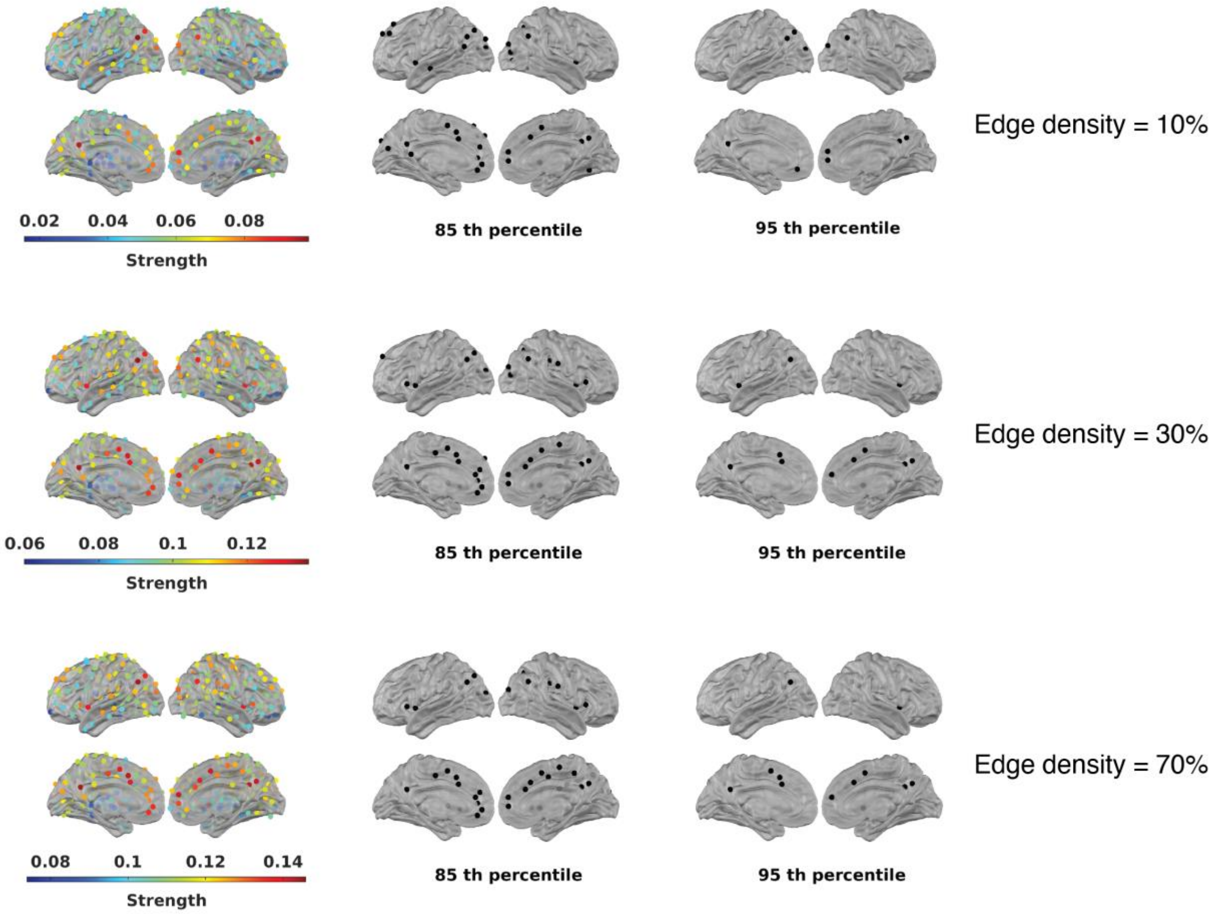


**Supplementary Figure. 5 Distribution of reference nodal strength and high strength hubs at different edge density thresholds.** Edge density 10%, 30%, 70%. ROIs with strength higher than the 85$^{th}$ percentile and 95$^{th}$ percentile are shown in the middle and right column.

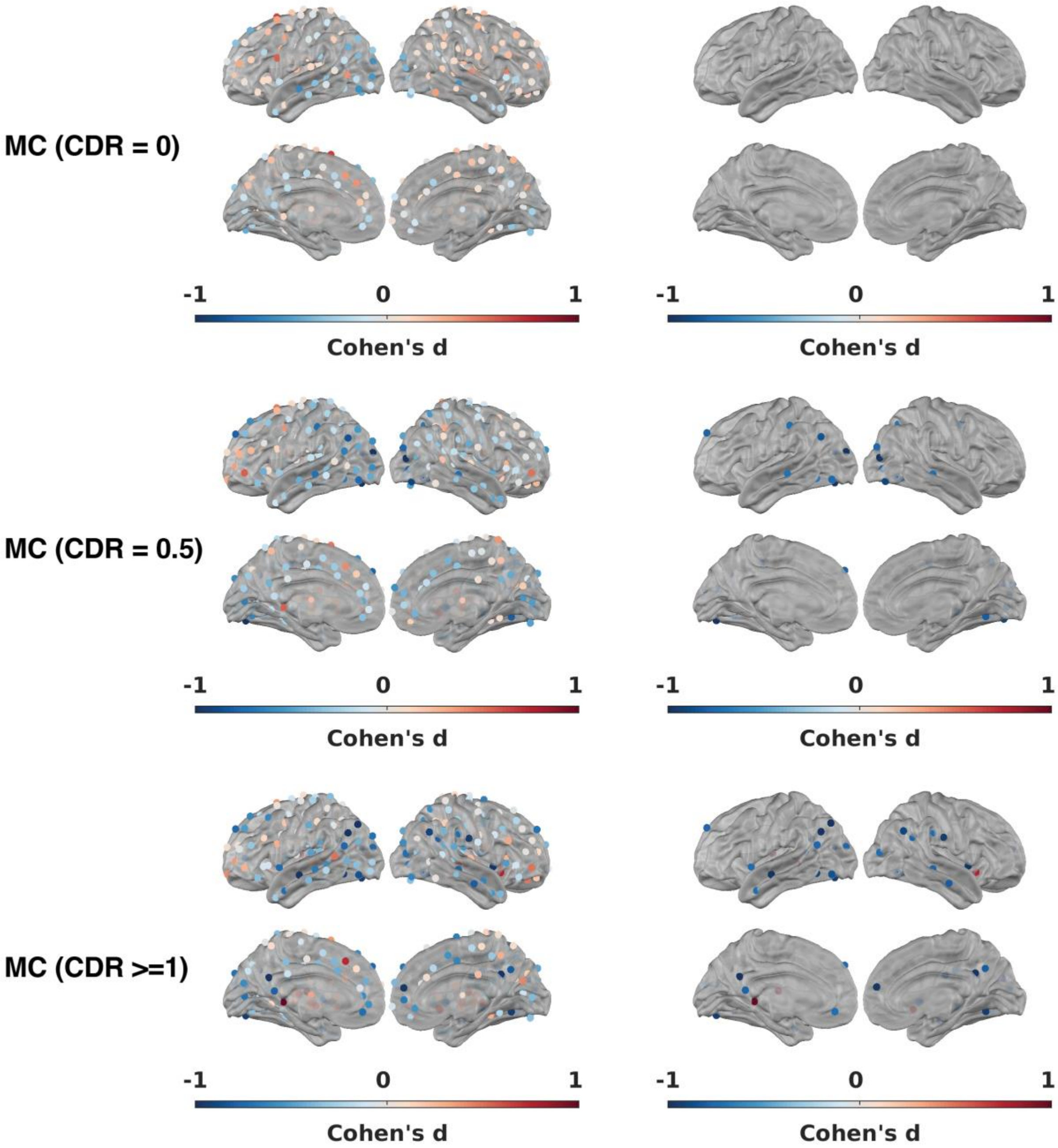


**Supplementary Figure. 6 Effect size of S difference from baseline and regions with a significant change from baseline (two-sample t-test, FDR<0.05) for mutation carriers.** Left column: Cohen's d for all ROIs. Right column: Cohen's d for ROIs passing an FDR<0.05 threshold.

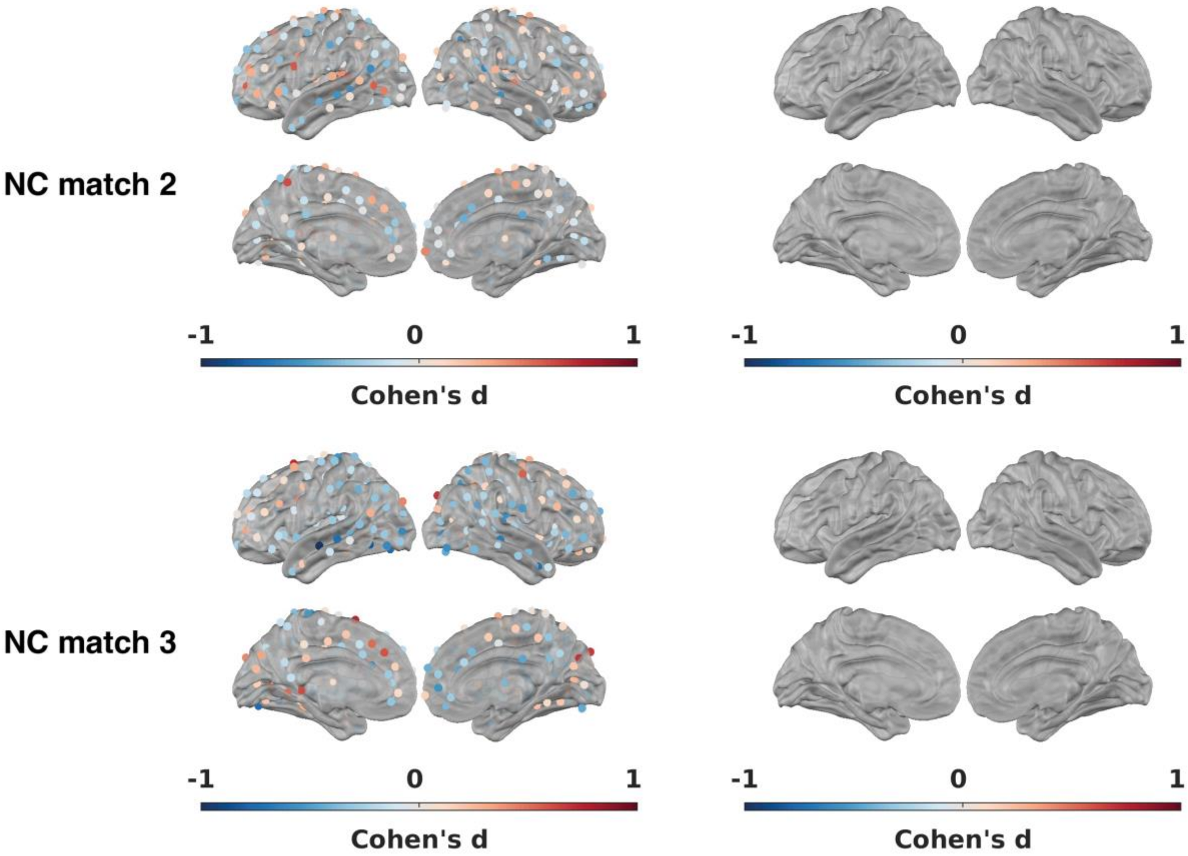


**Supplementary Figure 7. Effect size of S difference from baseline and regions with a significant change from baseline (two-sample t-test, FDR<0.05) for mutation non-carriers.** Left column: Cohen's d for all ROIs. Right column: Cohen's d for ROIs passing an FDR<0.05 threshold.

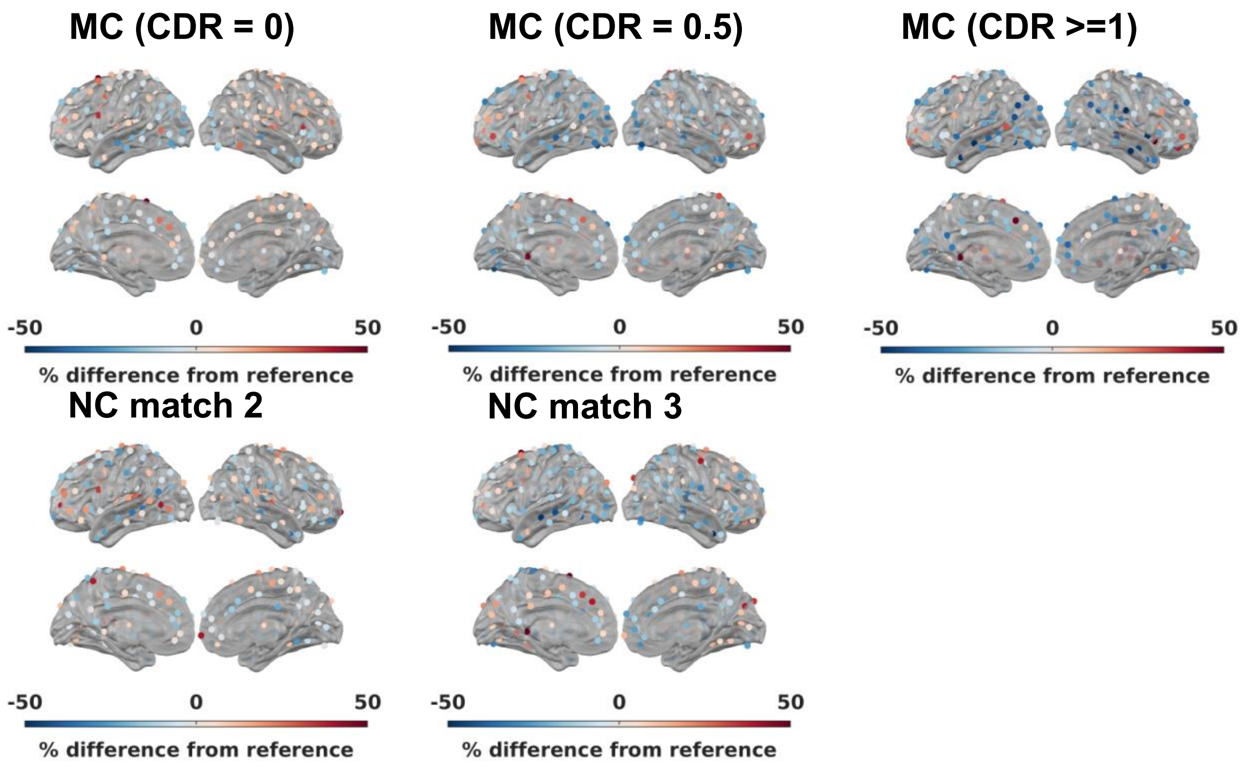


**Supplementary Figure 8. Visualization of % strength change on the brain. Edge density = 5%.**

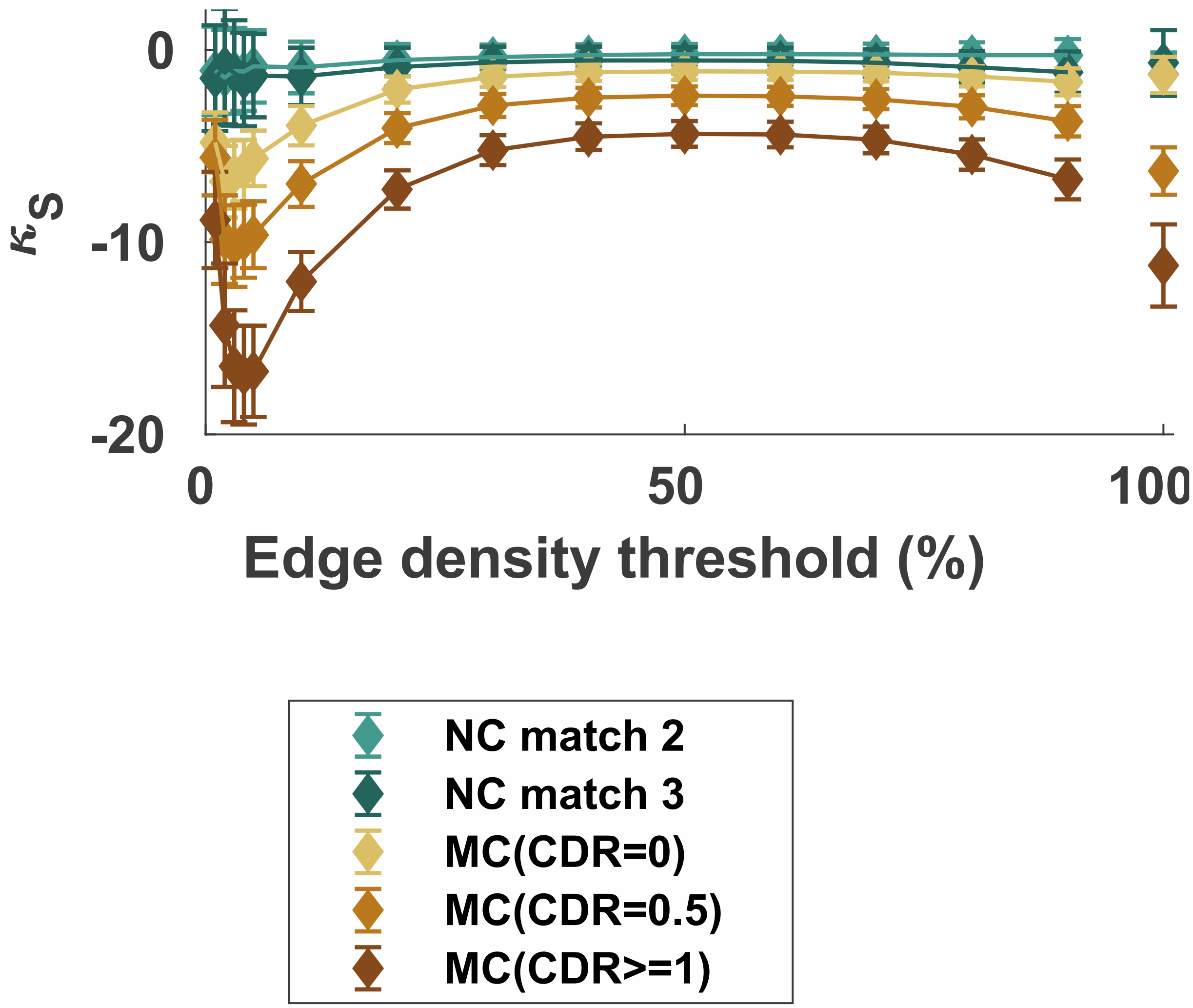


**Supplementary Figure 9. Hub disruption in strength at different edge densities**

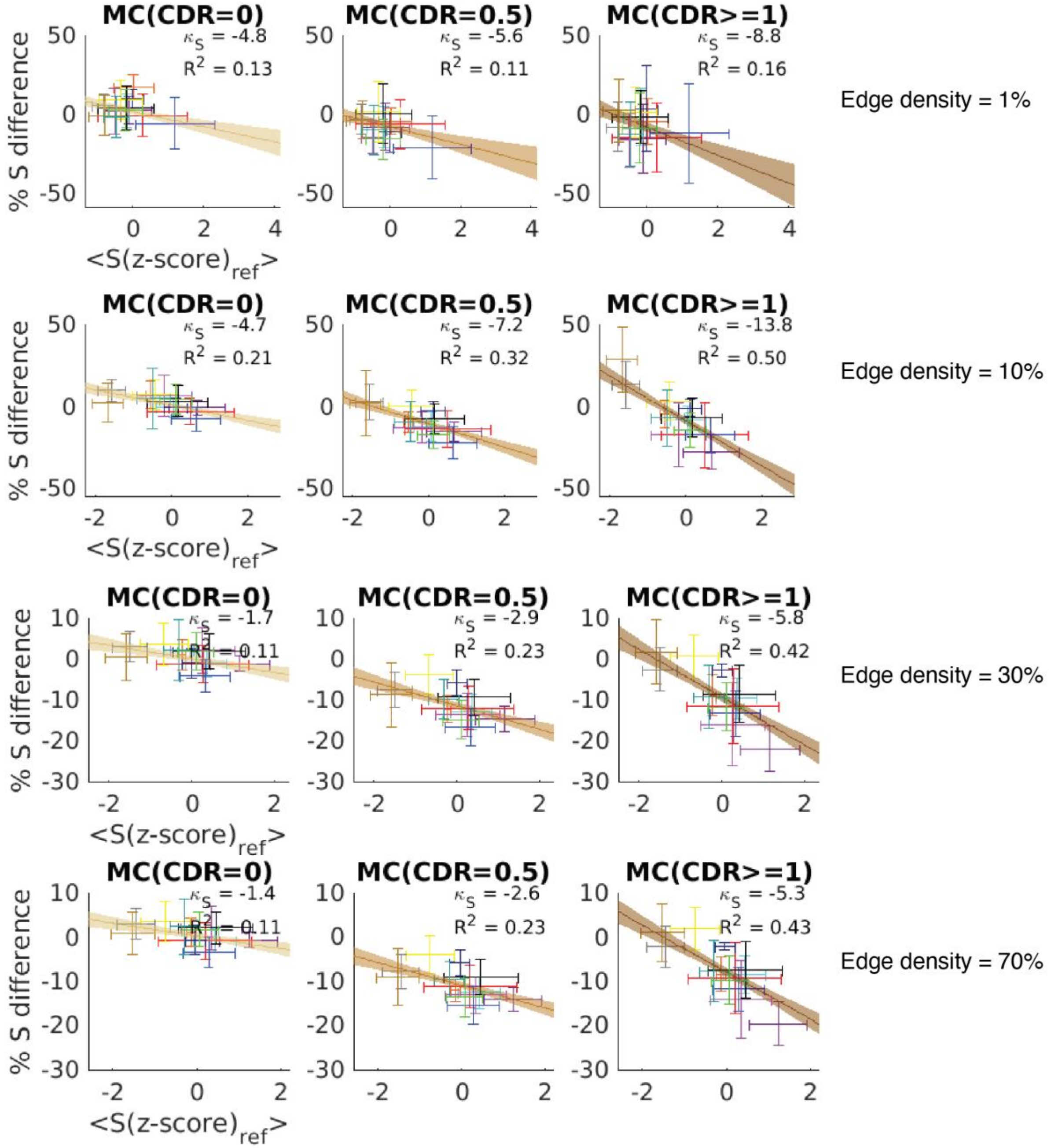


**Supplementary Figure 10. Network summary of %S difference from reference against S(z-score) in NC match 1.** Edge density = 1%, 10%, 30%, 70%. The cross hairs show the mean + SD of the %S difference from reference against S(Z-score) for individual networks across all ROIs in the network. Shaded areas. The networks are color-coded according to Figure 1A. 13 Networks: SMd, somatomotor dorsal; SMl, somatomotor lateral; CO, cingulo-opercular; AUD, auditory; DMN, default mode network; Mem, memory network; Vis, visual network; FPN, frontoparietal network; SN, salience network; BG, basal ganglia; Thal, thalamus; VAN, ventral attention network; DAN, dorsal attention network.

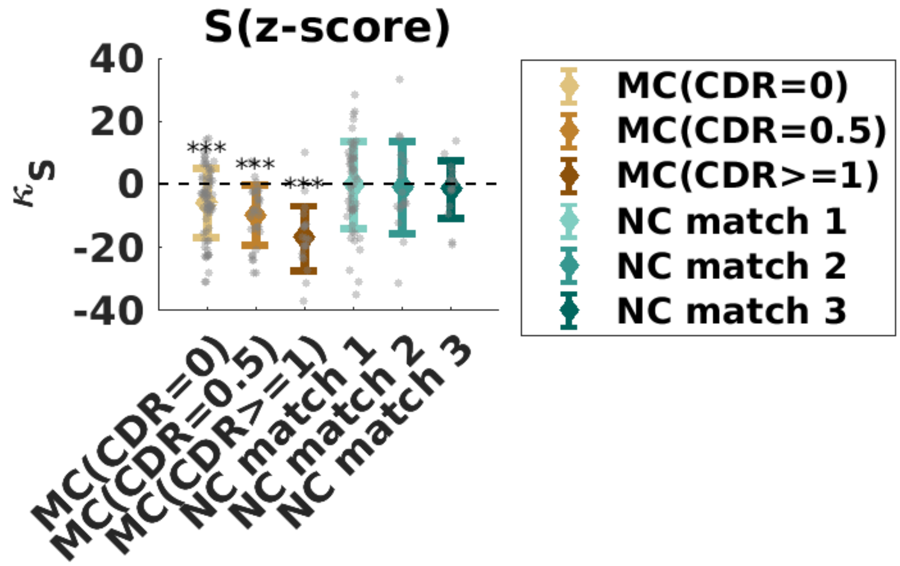


**Supplementary Figure 11. Comparing the hub disruption indices with zero.** The three mutation carrier groups have hub disruption indices with mean smaller than zero. The three non-carrier groups have hub disruption indices with mean not significantly different than zero. One-sample t-test with two-tails. Bonferroni-corrected p-values. * p<0.05, ** p<0.01, *** p<0.001.

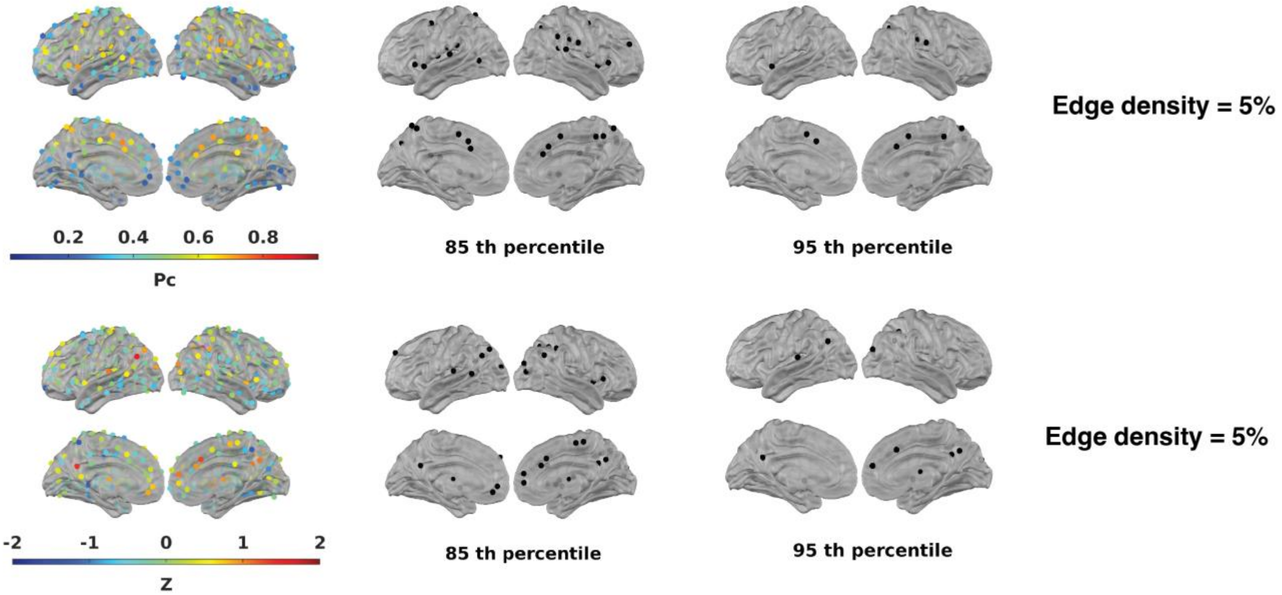


**Supplementary Figure 12. Participation coefficient (Pc) and within-module strength Z-score (Z) and the areas passing 85$^{th}$ percentile or 95$^{th}$ percentile thresholds.** Edge density = 5%.

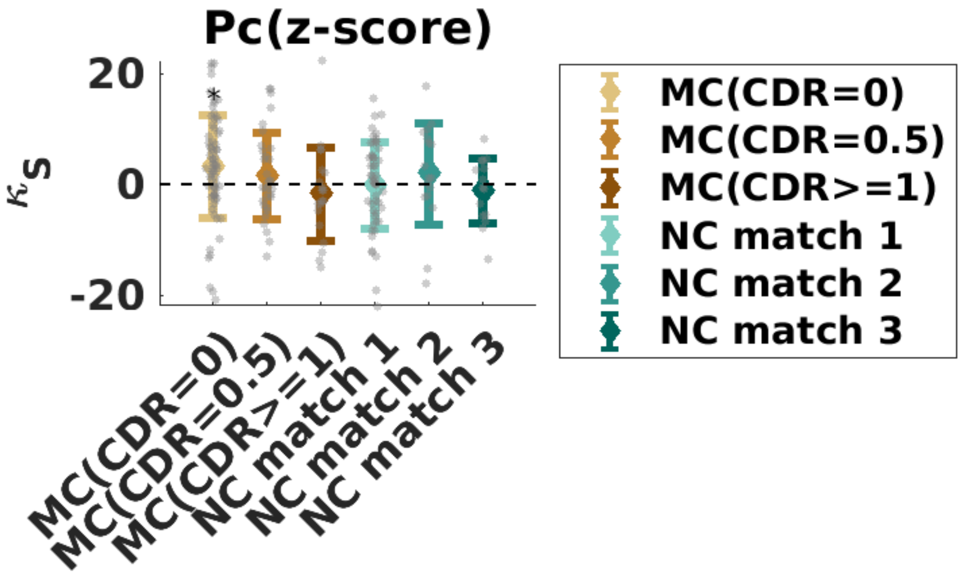


**Supplementary Figure 13. Individual hub disruption indices (with Pc as reference) compared to zero.** One-way ANOVA and post-hoc t-test with Bonferroni correction for six tests. *p<0.05, **p<0.01, ***p<0.001

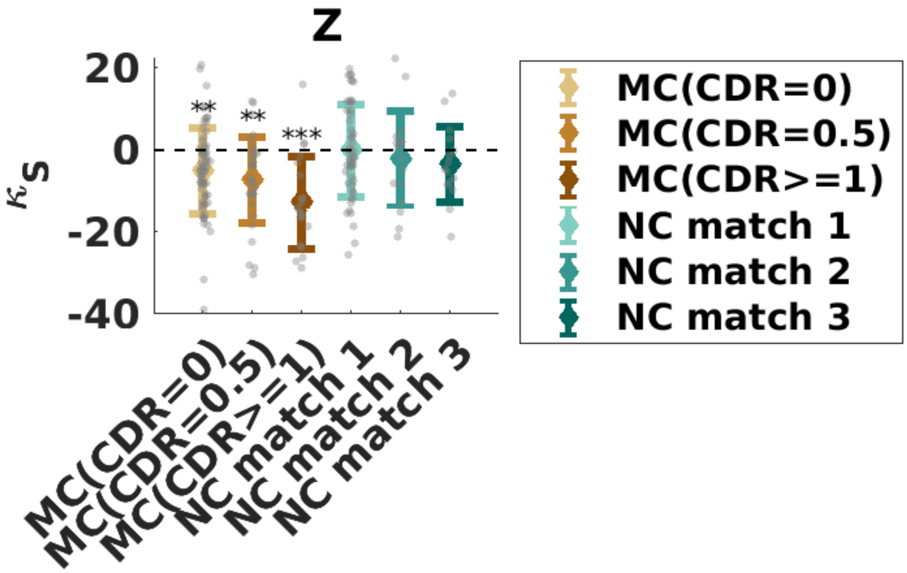


**Supplementary Figure 14. Individual hub disruption indices (with Z as reference) compared to zero.** One-way ANOVA and post-hoc t-test with Bonferroni correction for six tests. *p<0.05, **p<0.01, ***p<0.001

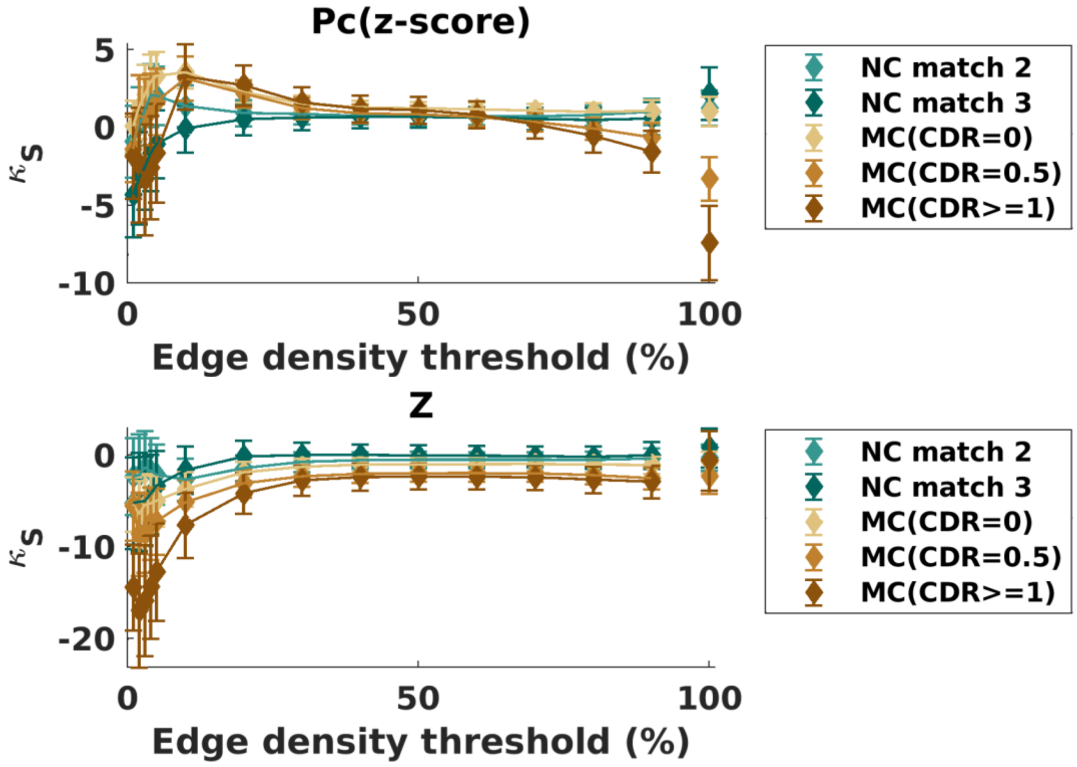


**Supplementary Figure 15. Change in strength in relation to (top) participation coefficient (Pc) and (bottom) within-module strength Z-score (Z) across thresholds.**

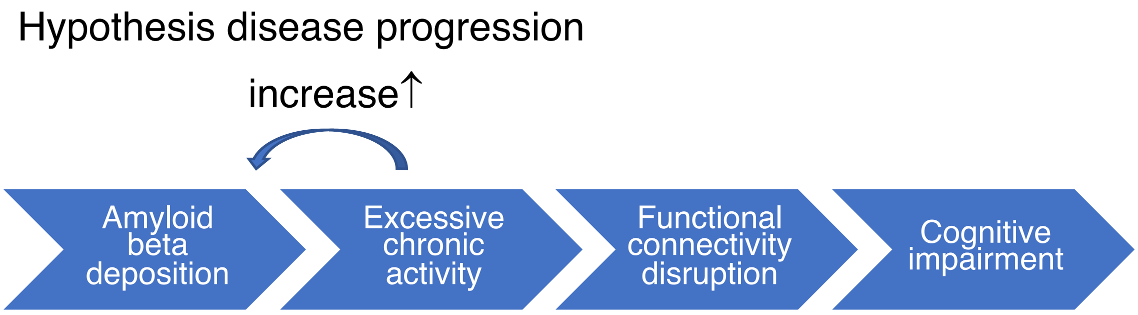


**Supplementary Figure 16. Hypothesis disease progression model**

### Appendix

#### A. Test for sufficiency of data amount to produce reliable group-level RSFC measures

Since the size of the three groups is different by a large number (N = 69, 32, 20 for MC CDR = 0, CDR = 0.5, CDR>=1, respectively), one concern is that the group difference is a result of the difference in the number of participants in each group. To address this possible confound, we calculated the reliability of mean functional connectivity, mean nodal strength, and the z-score of mean nodal strength for different numbers of subjects in the NC match 1 group following the guidance of previous literature (Laumann et al. 2015), and demonstrate that convergence of those measures is seen at ~5 participants and ~20 min of total low-motion data, much less than any of our groups (Supplementary Figure. 18). This suggested that the number of subjects is enough for for group-level RSFC measures.

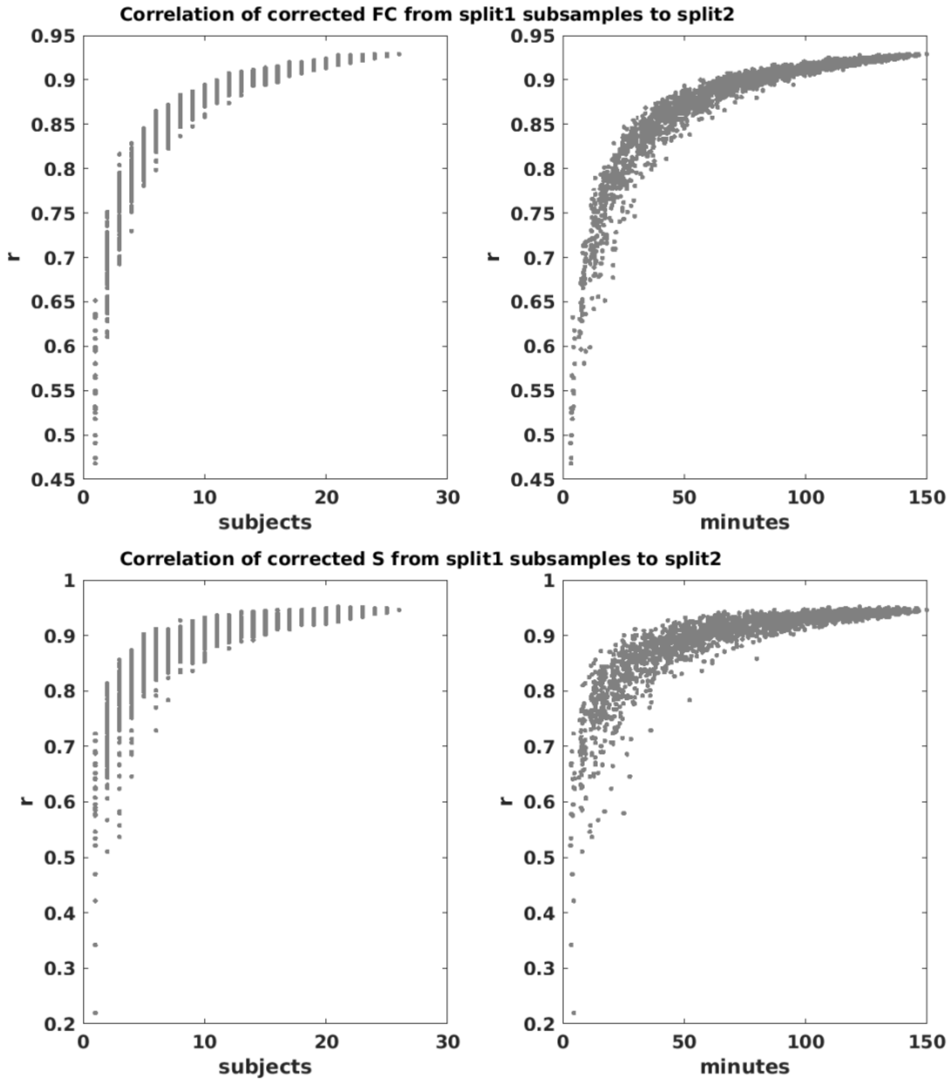


**Supplementary Figure 18 Mean functional connectivity (FC), mean nodal strength (S) and z-score of mean nodal strength (S(z-score)) similarity between subsamples of split 1 (26 subjects, 150 minutes) and that in split2 (26 subjects, 163 minutes) of the NC match 1 group (young non-carrier control). Edge density = 5%.** The similarity is measured with Pearson's r. This suggest that except for several outliers, the FC and nodal strength of subjects with ~4 minutes of low motion data have reasonable reliability (r~0.6), and reliability plateaued at ~5 subjects and ~20 minutes of low motion data, much lower than our group size.

### Appendix. B

Using the mean of the same young, healthy control (NC match 1) as a reference, we found a similar distribution of Pc and Z to previous literature (Power et al. 2013) (Figure4A; Supplementary Figure 12). We calculated the group-level hub disruption index with NC match 1 following the same process but with average Pc as the centrality measure (Figure 4B). We found that the reduction of functional connectivity is not disproportionally higher at connectors with high Pc (Table 2). This observation was also confirmed in the individual-level hub disruption index (Supplementary Figure 13), where a one-way ANOVAs showed no significant difference across the MC groups ($F (2,118) = 2.5$, $p=0.09$, Figure 4C) or the NC groups ($F (2, 81) =0.7$, $p=0.51$; Figure 4D).

The group-level hub disruption index with Z in NC match 1 as the reference (Figure 4E) showed a qualitatively similar pattern to the results using the NC match 1 average global connectivity strength as a reference (Figure 3): all MC groups had a significant negative hub disruption index while the NC groups did not (FDR-adjusted $p<0.05$) (Table 2). We found no significant differences in slopes between MC (CDR = 0.5) and MC (CDR = 0) ($F (1,488) = 0.9$, $p=0.33$), or between MC (CDR=0.5) and MC (CDR >=1) ($F (1,488) = 2.8$, $p=0.09$). However, there was a significant difference in slope between MC (CDR=0) compared to MC (CDR >=1) ($F (1,488) = 6.5$, $p=0.01$). We also conducted a one-sample t-test on the individual hub disruption index and obtained the same results (Supplementary Figure 14). All MC groups have hub disruption as indexed with a regression slope <0 (FDR-adjusted $p<0.05$): MC (CDR=0) ($M = -5.0$, $SD = 10.3$, Cohen’s $d = -0.5$, $t(68)=-4.0$, $p<0.001$); MC (CDR=0.5) ($M = -7.2$, $SD = 10.6$, Cohen’s $d = -0.7$, $t(31)=-3.9$, $p=0.001$); MC (CDR>=1) ($M = -12.7$, $SD = 11.3$, Cohen’s $d = -1.1$, $t(19)=-5.0$, $p<0.001$). A one-way ANOVA on the MC groups showed that there were significant differences in at least one of the pairs ($F (2,118) = 4.2$, $p=0.02$) (Figure 4F). A post-hoc t-test showed that MC (CDR=0) and MC (CDR>=1) groups had significantly different hub disruption index (FDR-adjusted $p = 0.01$). In contrast, a one-way ANOVA on the NC groups showed no significant difference in any of the pairs ($F (2, 81) =0.6$, $p=0.54$) (Figure 4G). Moreover, we validated our group-level results at different edge density thresholds to show that our results are consistent across thresholds (Supplementary Figure 15).

Wechsler, David. 1981. *WAIS-R Manual: Wechsler Adult Intelligence Scale-Revised*. Psychological Corporation.

———. 1987. *WMS-R: Wechsler Memory Scale--Revised : Manual*. Psychological Corporation.
